# Conditional GWAS of non-CG transposon methylation in *Arabidopsis thaliana* reveals major polymorphisms in five genes

**DOI:** 10.1101/2022.02.09.479810

**Authors:** Eriko Sasaki, Joanna Gunis, Ilka Reichardt-Gomez, Viktoria Nizhynska, Magnus Nordborg

## Abstract

Genome-wide association studies (GWAS) have revealed that the striking natural variation for DNA CHH-methylation (mCHH; H is A, T, or C) of transposons has oligogenic architecture involving major alleles at a handful of known methylation regulators. Here we use a conditional GWAS approach to show that CHG-methylation (mCHG) has a similar genetic architecture — once mCHH is statistically controlled for. We identify five key *trans*-regulators that appear to modulate mCHG levels, and show that they interact with a previously identified modifier of mCHH in regulating natural transposon mobilization.

## Introduction

Organisms have developed defense systems to protect the genome from transposable elements, which can act as ‘selfish genes’ and cause considerable damage (Chuong et al., 2017; Deniz et al., 2019). Cytosine DNA methylation is a major component of genome defense that is found in both mammals and plants, albeit with significant differences (Law and Jacobsen, 2010; Deniz et al., 2019). For instance, whereas mammals mostly have CG methylation (mCG), methylation in plants also occurs in the CHG and CHH contexts (H is A, T, or C). mCG in plants is known to be maintained through both mitosis and meiosis by *DNA METHYLTRANSFERASE 1* (*MET1; DNMT1* in humans) — in contrast to CHH methylation (mCHH), which is re-established after cell division by several pathways, including the RNA-directed DNA methylation (RdDM) pathway and the *CHROMOMETHYLASE 2* (*CMT2*) pathway (Kawashima and Berger, 2014; Matzke et al., 2015). Unlike mCG, which is stably inherited, mCHH behaves like a molecular phenotype and is strongly influenced by the environment, such as growth temperature (Dubin et al., 2015) and stress (Wibowo et al., 2016). CHG methylation (mCHG) falls somewhere between mCG and mCHH in the sense that it can be maintained via positive feedback between *CHROMOMETHYLASE3* (*CMT3*) and *KRYPTONITE* (*KYP*), which recognize dimethylation of histone 3 lysine 9 (H3K9me2) and mCHG, respectively (Lindroth et al., 2001; Jackson et al., 2002; Cao and Jacobsen, 2002; Du et al., 2015). Molecular mechanisms aside, the forces shaping variation in DNA methylation remain obscure, and the same is true for their biological significance (Riddle and Richards, 2002; Reinders et al., 2009; Becker et al., 2011; Takuno and Gaut, 2012; Meng et al., 2016; Wibowo et al., 2016; Johannes and Schmitz, 2019; Muyle and Gaut, 2019; Baduel and Colot, 2021). Previously, we demonstrated the existence of large-scale geographic clines for DNA methylation in *A. thaliana*, and used genome-wide association studies (GWAS) to show that mCHH on transposons is heavily influenced by major polymorphisms at *trans*-acting modifiers corresponding to known DNA methylation regulators in silencing pathways: *CMT2, NUCLEAR RNA POLYMERASE D1B* (*NRPE1*), *ARGONAUTE 1* (*AGO1*), and *AGO9* (Dubin et al. 2015, Kawakatsu et al 2016, Sasaki et al 2019). In particular, an allele of *NRPE1* (named *NRPE1’*) strongly affects mCHH levels in RdDM-targeted transposons. *NRPE1*, which is the largest subunit of RNA polymerase V, localizes to promoter regions of relatively young transposons (Zhong et al., 2012), and Baduel et al. (2021) recently showed that the *NRPE1’* allele has been associated with recent transposon mobilization, presumably through its effect on DNA methylation. All in all, these findings are suggestive of a highly variable genome defense system.

While both mCHG and mCG showed high heritability, GWAS yielded little in terms of significant associations. This might be because these “traits” are highly polygenic, or because they are at least partly transgenerationally inherited, and hence do not behave like standard phenotypes. In this paper we revisit mCHG variation and use a conditional GWAS approach to reveal that multiple major alleles at *trans*-acting regulators modify this type of methylation, and are also associated with transposon mobilization.

## Results

### mCHG is strongly correlated with mCHH

Our starting point is the observation that mCHG and mCHH levels on transposons are strongly correlated in the 1001 Epigenomes data set (Kawakatsu et al., 2016), especially for RdDM-targeted transposons (Fig. 1A; see Methods). Much of this variation is due to differences in the environment (including tissue, which can be viewed as a cellular environment), with flower tissue samples showing clear hyper-mCHG compared with leaf samples as expected (Feng et al., 2020; Gutzat et al., 2020). At the same time, variation across individuals is huge even when controlling for known tissue and environmental effects, as observed in the largest leaf sample data set (“Leaf SALK ambient temperature”; *n*=846).

**Fig 1.**
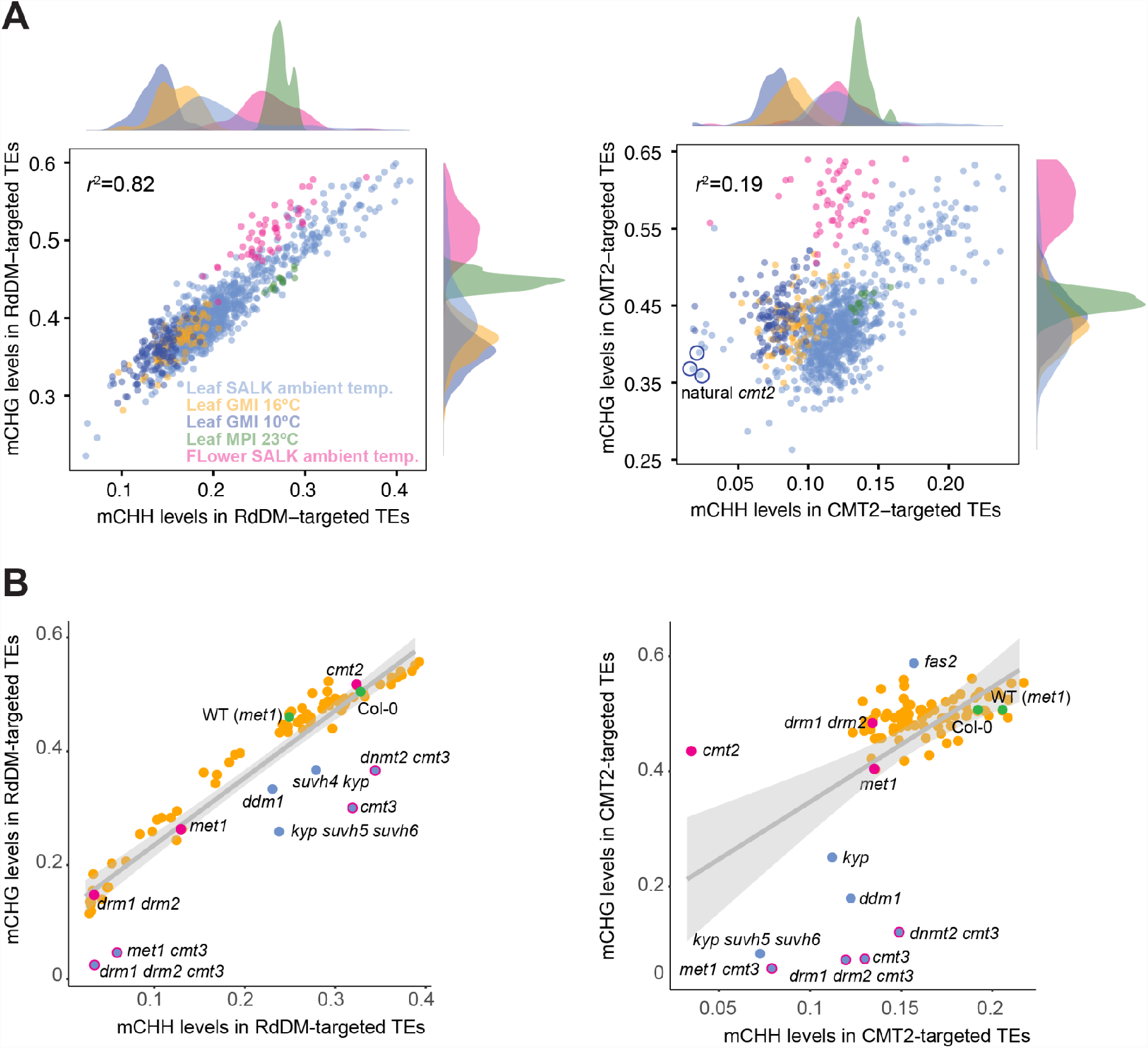
Covariance between mCHH and mCHG levels across individuals. **(A)** Correlation of genome-wide average mCHH and mCHG levels in RdDM-and CMT2-targeted transposons across 1028 natural inbred lines measured in different conditions (including 79 lines measured in more than one condition; see Kawakatsu et al., 2016) Colors correspond to environments/tissues. Plots on axes show marginal densities. Circled lines carry known natural null alleles of *CMT2*. **(B)** The same correlation as in A, but for 85 epigenetics-related loss-of-function mutants (Stroud et al., 2013). Gray areas indicate 99% confidence intervals around the linear regression lines. Green points denote “wildtypes” (see Stroud et al., 2013); magenta, major DNA methyltransferases. Blue points denote lines with a highly significant effect on mCHG.

Interestingly, the covariance between mCHH and mCHG showed the same pattern in data generated by knocking out known or potential DNA methylation regulators in the same genetic background (Fig. 1B) (Stroud et al., 2013). This demonstrates strong co-regulation of these types of methylation, in particular for RdDM-targeted transposons. Loss of RdDM regulation, such as in the double mutant of *DOMAINS REARRANGED METHYLASE (DRM) drm1 drm2*, causes loss of almost all mCHH and mCHG because mCHG in these regions is mainly established via siRNA (Chan et al., 2006). At the same time, the data demonstrate that some genetic perturbations can affect one type of methylation much more than the other. For example, knocking out *CMT3* affects mCHG levels much more than mCHH levels.

Based on these observations, it is clear that a substantial fraction of variation in non-CG methylation is due to factors that affect both mCHH and mCHG. Without replicate measurements, it is difficult to say what fraction of these factors is genetic and what is environmental, but, regardless of this, we hypothesized that the substantial covariance could reduce power of GWAS for either mCHH or mCHG (when using a standard univariate model), and that an analysis accounting for this covariance might perform better (Korte et al., 2012; Stegle et al., 2012; Stephens, 2013). In essence, we sought to simplify a complex trait by breaking it into constituent parts (Nilsson-Ehle, 1909; East, 1910; Lande and Arnold, 1983). This insight is the basis for this paper.

### Conditional GWAS reveals new associations

Multi-trait GWAS can be carried out using transformations (e.g., the difference or ratio between traits), conditional approaches in which one or more traits are accounted for as covariates, or full multivariate models (Korte et al., 2012; Stegle et al., 2012; Stephens, 2013). We tried several different approaches, including a multivariate GWAS, but this paper will focus on results from a conditional GWAS of mCHG with mCHH as a covariate (denoted mCHG_|mCHH_) because this gave the clearest results. Other approaches produced consistent results, and we will briefly discuss them later.

Figure 2 contrasts the performance of univariate and conditional GWAS. As previously noted, univariate GWAS of mCHG does not yield any significant associations despite moderate SNP-heritability (37% and 38% for RdDM-and CMT2-targeted transposons, respectively). As expected given the very strong correlation between mCHH and mCHG on RdDM-targeted transposons, the strong associations at *AGO1* and *NRPE1* found in univariate GWAS of mCHH (Kawakatsu et al., 2016) give rise to associations for mCHG as well, but they are not genome-wide significant (*AGO1* Chr1:17895231, -log_10_*p*-value=6.51 and *NRPE1* Chr2:16714815, -log_10_*p*-value=5.23). The previously identified *CMT2*-association with mCHH on CMT2-targeted transposons (Dubin et al., 2015; Kawakatsu et al., 2016; Sasaki et al., 2019; Shen et al., 2014) is not apparent, consistent with the much weaker correlation between mCHH and mCHG on these transposons.

**Fig 2.**
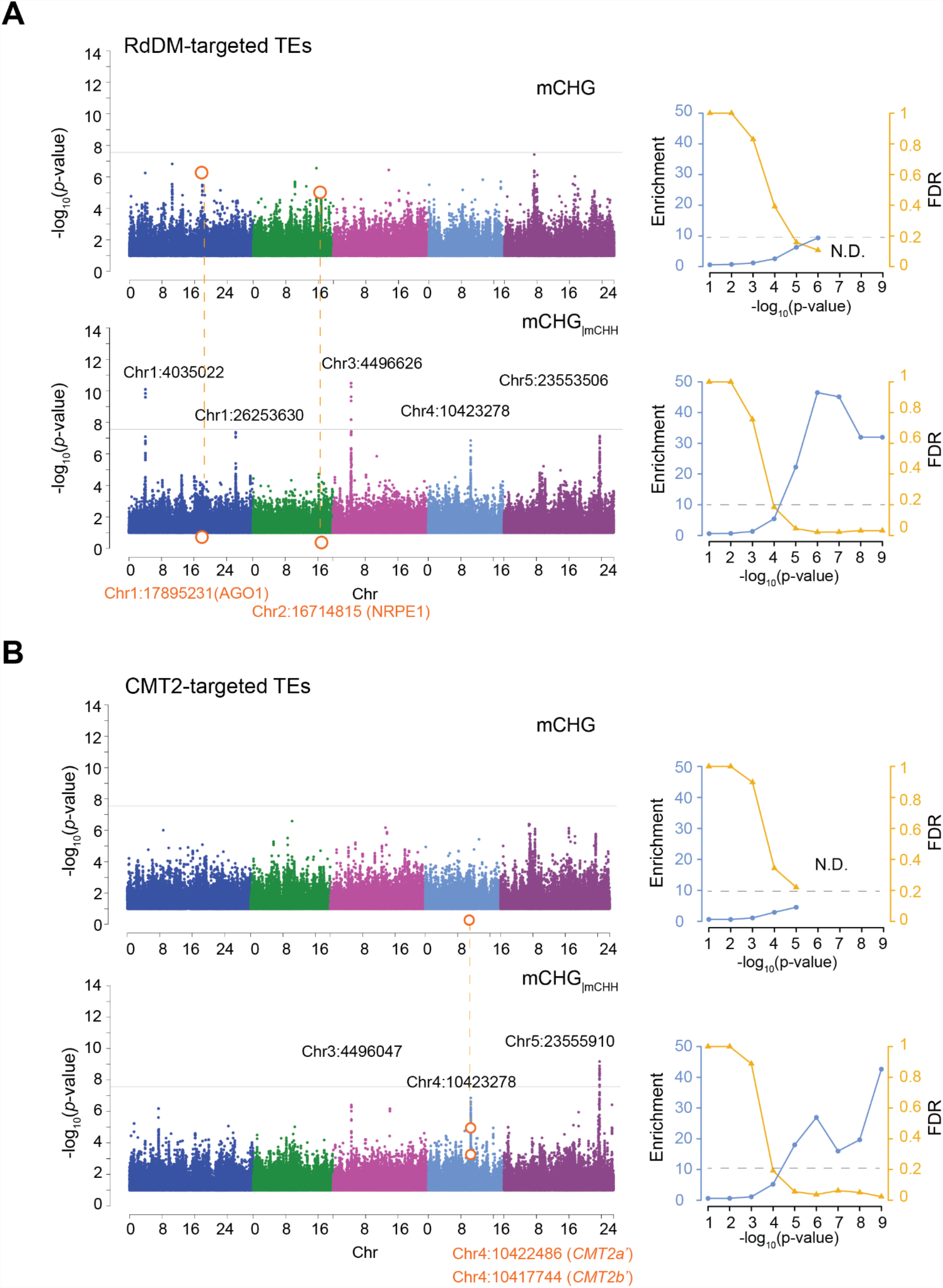
Comparison of univariate and conditional GWAS of mCHG. The analysis was done separately for (**A)** RdDM-targeted and (**B**) CMT2-targeted transposons, using the 774 lines in the global panel from the 1001 Epigenomes Project (“SALK leaf in ambient temperature”; see Fig. 1). For each case, the upper Manhattan plot shows univariate GWAS of mCHG methylation and the lower GWAS of mCHG controlling for mCHH. Horizontal gray lines show genome-wide significance (p=0.05 after Bonferroni-correction). The line plots show enrichment of *a priori* genes and FDR (see text), with horizontal dashed lines indicating an FDR of 20%.

The contrast between these univariate GWAS results and the conditional analysis is striking. GWAS for mCHG while controlling for mCHH (mCHG_|mCHH_) revealed five clear peaks for RdDM-targeted transposons, three of which were also found for CMT2-targeted transposons. The peaks are above or near genome-wide significance using a conservative threshold (Figs. 2, S1; Table S1). As will be discussed further below, these associations jointly explain 30.5% and 14.8% of mCHG_|mCHH_ in RdDM-and CMT2-targeted transposons, respectively. The previously mentioned *AGO1* and *NRPE1* associations disappear completely, consistent with their being strongly correlated with mCHH, and hence controlled for. The improved performance of conditional GWAS is also evident from a massive enrichment of associations near *a priori* candidates (Atwell et al., 2010). With conditional GWAS, we observed an up to 45-fold excess of associations near annotated epigenetic modifiers (Kawakatsu et al., 2016; Sasaki et al., 2019), compared to an 4-to 10-fold excess for the univariate analyses (Fig. 2).

The enrichment for *a priori* candidates also allows us to estimate a False Discovery Rate (FDR) for this set of genes (Atwell et al., 2010). Four of the five major associations can be identified with *a priori* candidates at very low FDRs (e.g. FDR < 0.05 using a significance threshold of -log_10_*p*-value > 6; see Fig. 2), but it is notable that FDR is low even for associations that are nowhere near genome-wide significance. This is not true for the univariate analyses. For example, at an FDR of 20%, univariate GWAS of mCHG identifies only *AGO1* and *NRPE1* for RdDM-targeted transposons, and nothing for CMT2-targeted transposons (Table S2), whereas conditional GWAS of mCHG_|mCHH_ generates a long list of known regulators of mCHG (Table S2). For RdDM-targeted transposons, we find four *a priori* genes above or near genome-wide significance: the previously mentioned *CMT2* and *CMT3*, plus *MIR823a*, which encodes a microRNA targeting *CMT3* (Rajagopalan et al., 2006; Papareddy et al., 2021), and *MULTICOPY SUPPRESSOR OF IRA1* (*MSI1*), likely a component of chromatin assembly factor (CAF-1) responsible for proper heterochromatin formation with *FASCIATA1* (*FAS1*) and *FASCIATA2* (*FAS2*) (Hennig et al., 2003, 2005). At lower significance levels, we find *REPRESSOR OF SILENCING 3* (*ROS3*) and *DNA METHYLTRANSFERASE 2* (*DNMT2*). *ROS3* is a DNA demethylase that interacts with *REPRESSOR OF SILENCING 1* (*ROS1*) (Zheng et al., 2008) and *DNMT2* is associated with histone deacetylation (Song et al., 2010) and RNA methyltransferase activity (Goll et al. 2006) (and could therefore be a false positive since the effects on DNA methylation is under debate; see Goll et al. 2006; Song et al. 2010). For CMT2-targeted transposons, the list of associated *a priori* candidates includes four genes also associated with RdDM-targeted transposons, namely *MIR823A, CMT2, ROS3*, and *MSI1*, but there are also two specific genes, *FAS1* and *DECREASED DNA METHYLATION 1* (*DDM1*), a chromatin-remodeler responsible for heterochromatin formation (Soppe et al., 2002; Osakabe et al., 2021). As shown in Fig. 1B, *ddm1* strongly reduces mCHG, and *fas2*, a functionally redundant *FAS1* homolog, increases CMT2-targeted mCHG in a mCHH-independent manner (Stroud et al., 2013).

### Genes underlying major associations

#### On searching for causal genes

Identifying causal genes and mechanisms from GWAS results is notoriously difficult (Gallagher and Chen-Plotkin, 2018). Peaks of association often cover multiple genes — this is certainly true in the gene-dense genome of *A. thaliana* (Atwell et al., 2010) — and functional annotation is of limited use for complex traits with unknown genetic basis. We are in a stronger position, however, because, just like in our previous work on mCHH (Sasaki et al., 2019), the significant GWAS peaks (Fig. 2) are narrow, and clearly pinpoint genes *a priori* known to be specifically involved in well-defined molecular phenotypes (Table 1, Fig. 3). Hence we will discuss these candidates, and the indirect evidence supporting their causal role.

**Table 1.** Significant conditional GWAS hits for mCHG_|mCHH_*.

| SNPs | $-\log_{10}$<br>(*p*-value) | MAC | Distance from<br>*a priori* gene (bp) | Genes within 15K bp* |
| --- | --- | --- | --- | --- |
| *mCHG<sub>mCHH</sub> in RdDM-targeted transposons* |  |  |  |  |
| Chr1:4035022 | 10.1 | 77 | NA | AT1G11930, AT1G11940, AT1G11950 (JMJ26), AT1G11960 |
| Chr1:26253630 | 7.38 | 290 | 50 | **AT1G69770 (CMT3)** |
| Chr3:4496626 | 10.48 | 207 | 196 | AT3G13700, AT3G13710 (*PRA1.F4*), AT3G13720 (*PRA1.F3*), **AT3G13724 (MIR823A)**, AT3G13730 (*SYP90D1*) |
| Chr4:10423278 | 6.85 | 301 | 2069 | **AT4G19020 (CMT2)**, AT4G19030 (*NLM1*), AT4G19035 (*LCR7*) |
| Chr5:23553506 | 7.14 | 99 | 2506 | AT5G58200 (*ECT10*), AT5G58200, AT5G58210, AT5G58220 (*TTL*), **AT5G58230 (MSI1)**, AT5G58240 (*FHIT*) |
| *mCHG<sub>mCHH</sub> in CMT2-targeted transposons* |  |  |  |  |
| Chr3:4496047 | 6.42 | 210 | 775 | AT3G13700, AT3G13710 (*PRA1.F4*), AT3G13720 (*PRA1.F3*), **AT3G13724 (MIR823A)**, AT3G13730 (*SYP90D1*) |
| Chr4:10423278 | 6.86 | 301 | 2069 | **AT4G19020 (CMT2)**, AT4G19030 (*NLM1*), AT4G19035 (*LCR7*) |
| Chr5:23555910 | 9.17 | 64 | 102 | AT5G58200, AT5G58210, AT5G58220 (*TTL*), **AT5G58230 (MSI1)**, AT5G58240 (*FHIT*), AT5G58250, AT5G58260 (*NdhN*) |

\* Bold denotes *a priori* epigenetic regulators

**Fig 3.**
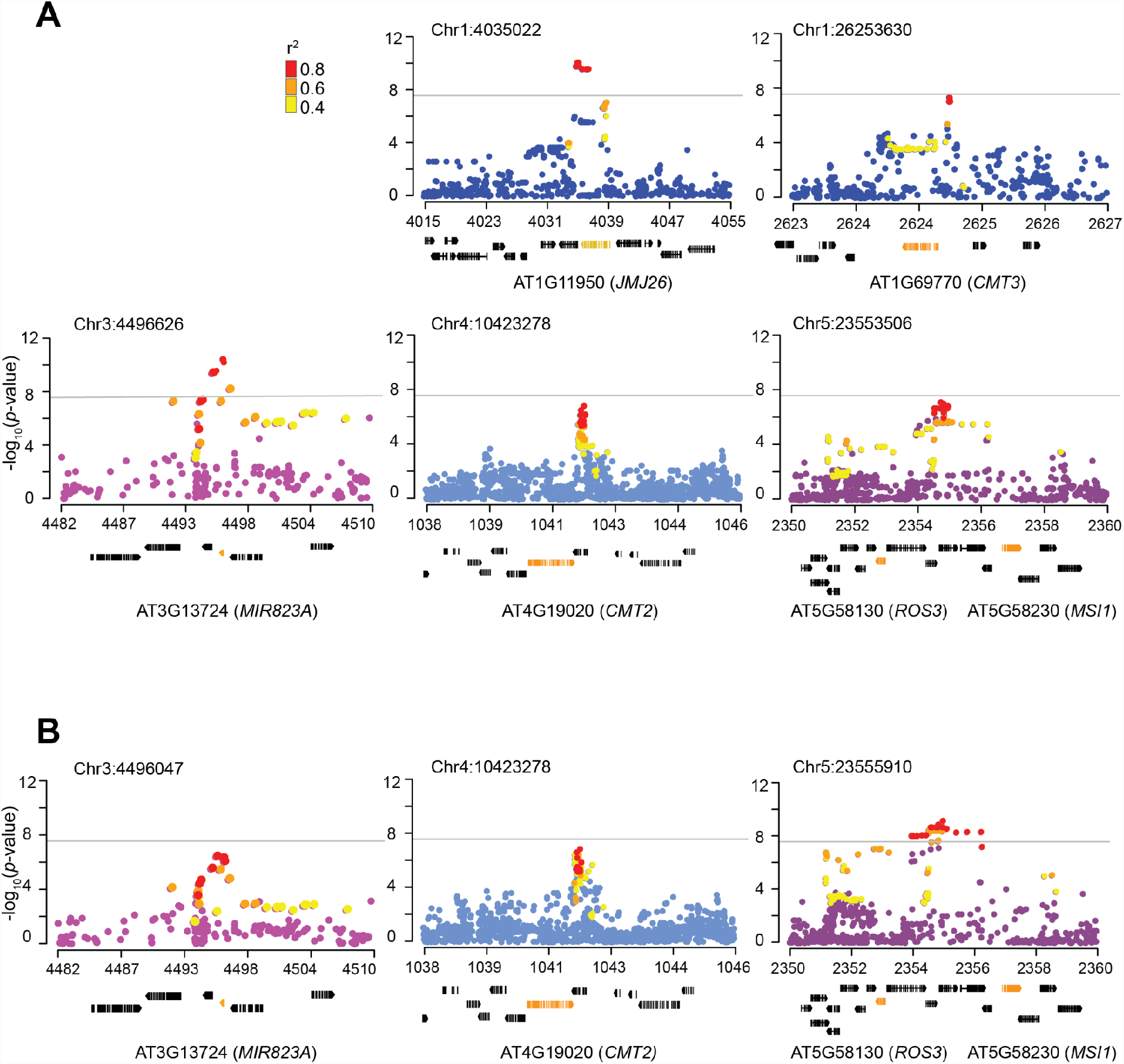
Candidate loci underlying mCHG_|mCHH_variation. Zoomed-in Manhattan plots aroundconditional GWAS peaks in Fig. 2 for **(A)** RdDM-and **(B)** CMT2-targeted transposons. Gene models below the plots show the candidate genes in orange (see Table 1). Colored dots represent SNPs in strong linkage disequilibrium with top SNPs.

#### Multi-layered direct *CMT3* regulation affects mCHG variation

Based on mutant phenotypes, *CMT3* is a strong *a priori* candidate for regulating mCHG (Fig. 1). Consistent with this, one of the five major associations from GWAS of mCHG_|mCHH_ pinpoints *CMT3* and another *MIR823a*, a gene encoding a microRNA that directly down-regulates *CMT3* by cleaving its transcripts during early embryogenesis (Papareddy et al., 2021). The most significant peak for mCHG_|mCHH_ on RdDM-targeted transposons was located 196 bp downstream of *MIR823a*, while the peak corresponding to *CMT3* was located 50 bp upstream of the gene (Fig. 4A, Table 1). GWAS for CMT2-targeted transposons also pinpointed *MIR823a*, although the association is weaker, and the most significant SNP further away (Fig. 3B, Table 1). This association was also recently found by Hüther et al. (2022) using GWAS for unconditional mCHG levels of individual transposons.

**Fig 4.**
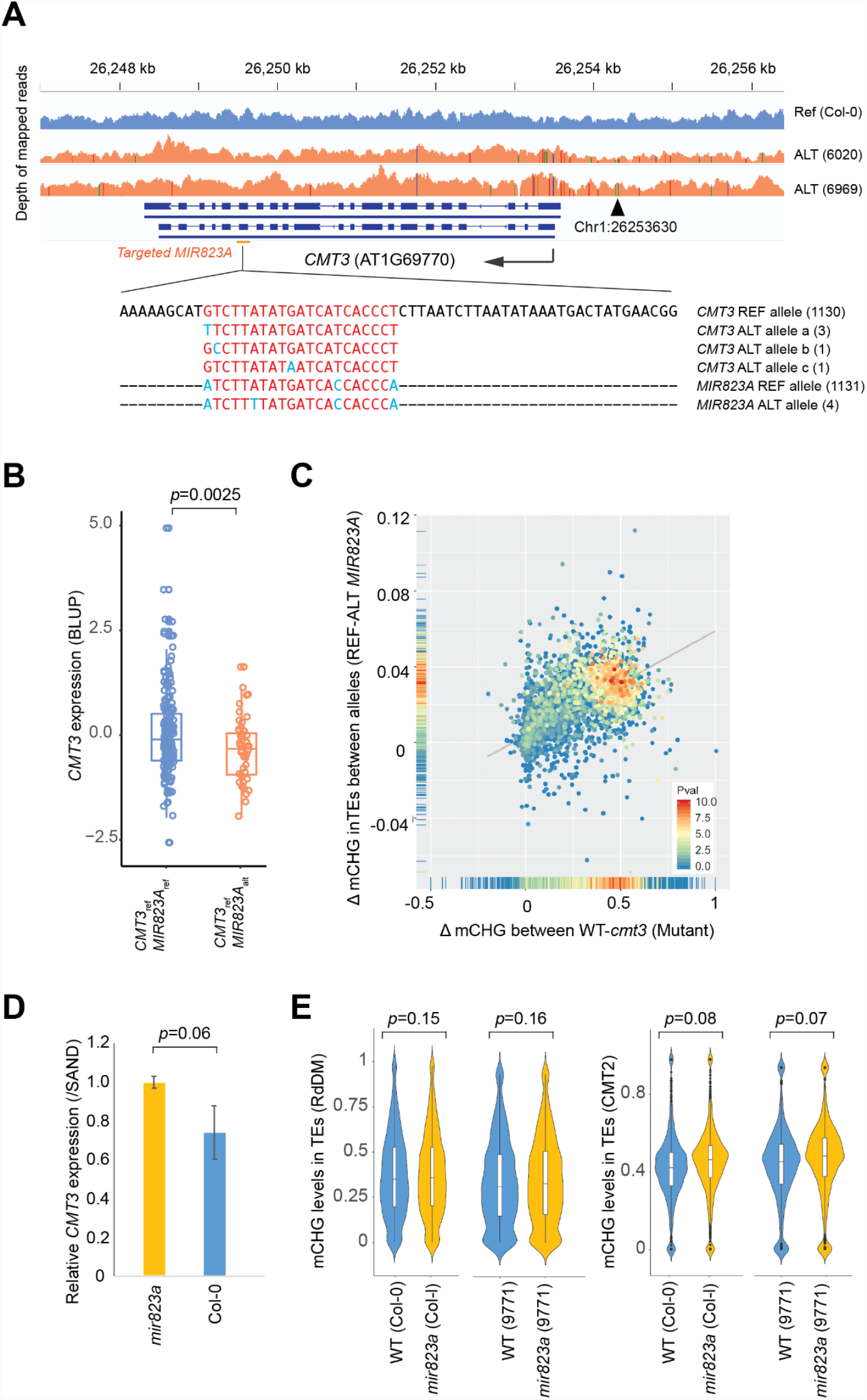
Effects of *MIR823A* on mCHG levels. **(A)** Polymorphisms in the miRNA region of *MIR823A* and the target in *CMT3* in 1135 natural lines. Differences from the *CMT3* reference sequence are shown in light blue. Haplotype counts are shown in parentheses. **(B)** Estimated effects of *MIR823A* alleles on *CMT3* expression in natural lines with individual values (Welch’s t-test, two-tailed, *CMT3* expression was collected from transcriptome data published in Kawakatsu et al., 2016). **(C)** Scatter plot comparing the average allelic effect of the *MIR823A* polymorphism with the effect of *cmt3* in the Col-0 background. Dots represent individual transposons, and the colors show the significance of the allelic effects as -log_10_p-value in GWAS. **(D)** *CMT3* expression in early embryos with standard errors in WT (Col-0) and *mir823a*. Significance of the difference in mean tested using Welch’s t-test (*n*=3, one-tailed). **(E)** The distribution of mCHG levels in RdDM and CMT2-targeted transposons, comparing knock-outs of *MIR823A* to wildtype in different backgrounds.

The *MIR823A* polymorphism appears to almost exclusively affect mCHG (Figs. S2, S3), primarily targeting the same transposons as a *CMT3* knock-out, as expected if the former directly regulates the latter (Fig. 4C). Consistent with this interpretation, lines carrying the non-reference *MIR823A* allele and a *CMT3* reference allele showed lower *CMT3* expression as well as lower mCHG (Fig. 4B). The specificity of the phenotypic effects and known regulatory mechanism provides strong evidence for a direct causal role for these genes.

Knock-outs of *MIR823A* in several backgrounds affected *CMT3* expression and mCHG in a manner consistent with this regulatory model (Papareddy et al., 2021), although the effects on methylation were very weak (Fig. 4D, E). The natural polymorphism almost certainly does not involve a loss of function, as there is no common polymorphism in the 21-nt mature miRNA region of *MIR823A*, nor in the target region of *CMT3* (Fig. 4A). There are also no other non-synonymous polymorphisms in *CMT3* significantly associated with the different alleles.

#### Further evidence for allelic heterogeneity at *CMT2*

In addition to *de novo* establishment of methylation in heterochromatic regions, *CMT2* plays a role in mCHG maintenance through regulation of H3K9 methylation via a self-reinforcing loop (Du et al., 2015; Stroud et al., 2014). Previous work has identified two common natural alleles, *CMT2a’* and *CMT2b’*, affecting mCHH (Dubin et al., 2015; Sasaki et al., 2019; Kawakatsu et al., 2016), but neither appears to affect mCHG (Figs S2, S3, S5). Here we identify a new association, 2 kb downstream of *CMT2* (Fig. 3), that does appear to affect mCHG (on both RdDM-and CMT2-targeted transposons) (Fig S5). The top SNP is only weakly correlated with *CMT2a’* and *CMT2b’* (*r*^*2*^_*CMT2b*’_=0.37; *r*^*2*^_*CMT2a’*_=0.24), but caution is needed when interpreting associations in regions that apparently harbor multiple causal polymorphisms (Sasaki et al., 2021).

### A complex association on chromosome 5 includes two *a priori* genes

The major peak on chromosome 5 was associated with mCHG_|mCHH_ on both RdDM-and CMT2-targeted transposons, although the shape of the peak differs slightly. The strongest association was found for CMT2-targeted transposons, and is located 102 bp upstream of a known epigenetic regulator, *MSI1*, but substantial linkage disequilibrium extends over a 30 kb region which also includes another *a priori* gene, *ROS3* (Figs 3, S6A).

The loss-of function mutant *ros3* does not show altered mCHG in leaves (Fig 1B; Stroud et al., 2013). Loss of *MSI1* causes embryonic lethality (Guitton et al., 2004), but the heterozygous mutant *msi1-2* does not show a significant effect on mCHG levels in leaves either. However, *MSI1* is required to control DNA methylation via repression of *MET1*, and a loss of *FAS2* in CAF-1 induces mCHG hypermethylation (Fig 1B) (Stroud et al., 2013; Jullien et al., 2008), so *MSI1* remains a strong candidate. Furthermore, the strongly associated SNP at Chr5:23555910 could explain almost all phenotypic variation associated with the chromosome 5 peak (Fig S6B), making *MSI1* the top candidate, although further experiments are clearly needed.

#### A jmjC gene is a novel modifier of mCHG in RdDM-targeted transposons

The final peak (Chr1:4035022; -log_10_*p-*value=10.1) in our study did not pinpoint any *a priori* gene. However, it is highly significant and narrowly centered on the coding region of a jmjC gene, *JUMONJI26* (*JMJ26*) (Qian et al., 2015). *JMJ26* is a close homolog of *JMJ25*, also known as *INCREASE IN BONSAI METHYLATION 1* (*IBM1*), a histone H3K9m demethylase targeting genic regions in a *KYP/SUVH4-*and *CMT3*-dependent manner (Saze et al., 2008; Inagaki et al., 2010). In contrast to *IBM1*, the function of *JMJ26* has barely been studied. Among the most significant associations were two non-synonymous SNPs (Chr1:4035683, 4035690) in the conserved JMJ domain, suggesting allelic effects on the enzymatic activity (Figs 5, S7).

**Fig 5.**
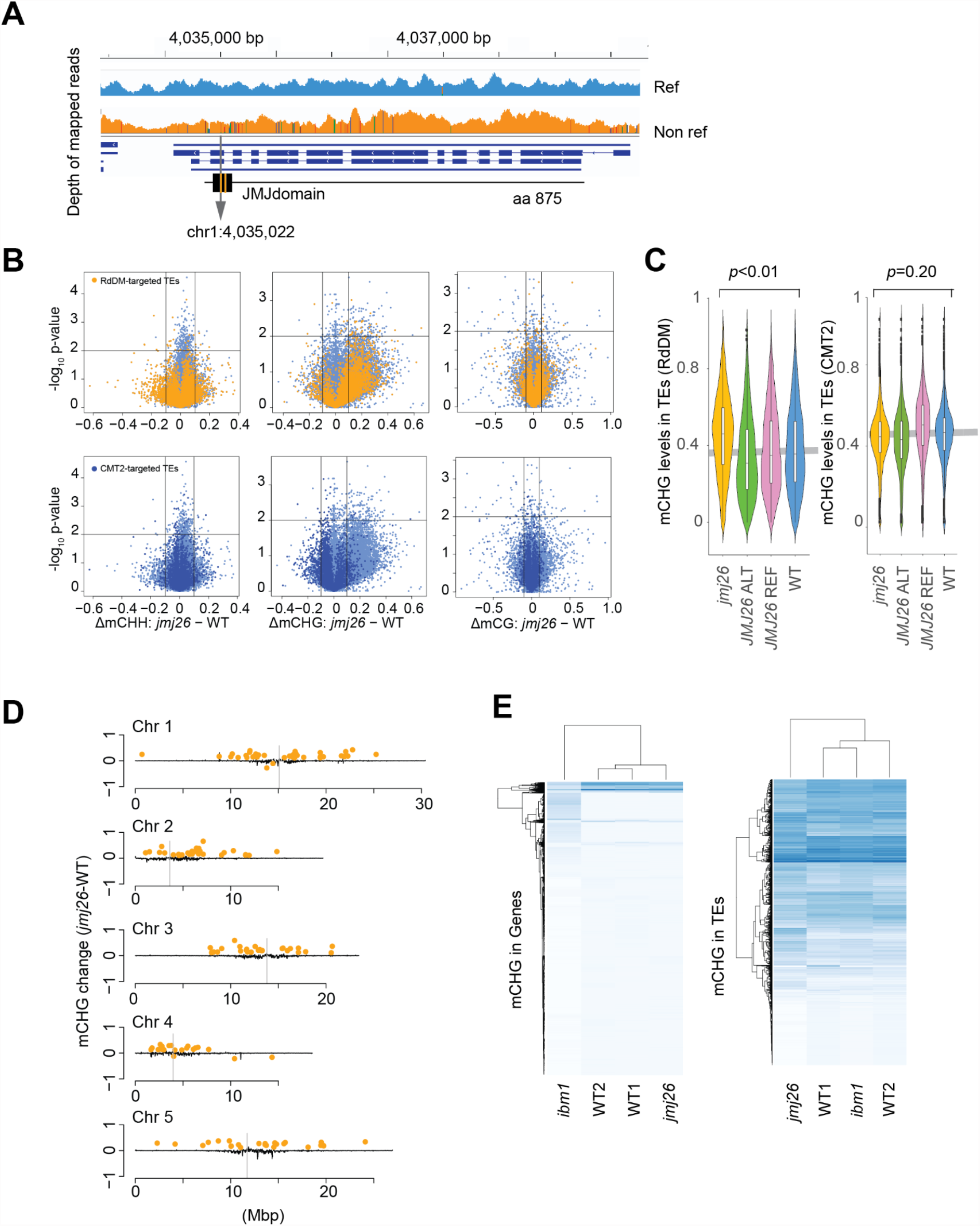
Effects of *JMJ26* on mCHG levels. **(A)** Read density and SNP across the gene model. **(B)** Volcano plots showing the effects on DNA methylation levels of *jmj26* on RdDM-and CMT2-targeted transposons. **(C)** Violin plots showing distribution of mCHG levels of RdDM-and CMT2-targeted transposons in WT (Col-0), *jmj26*, and two lines with *JMJ26*_ref_ or *JMJ26*_alt_ transgenes in Col-0 *jmj26* background. The gray horizontal line shows the mean value of WT. **(D)** The genome-wide distribution of transposons affected by *JMJ26*. Black lines show differential mCHG levels of *jmj26* from Col-0 in 30 kbp sliding windows. Orange dots show transposons with significantly changed mCHG levels (*n*=2, *p*<0.01 by two-tailed Welch’s *t*-test & differential mCHG > 0.1 or < -0.1). **(E)** Heat map comparing *JMJ26* and *IBM1* targets. Rows correspond to standardized mCHG levels in each gene or transposon for *jmj26, ibm1*, and WTs.

To explore the function of *JMJ26*, we measured mCHG levels of a loss-of-function mutant, *jmj26* (Fig. 5). Unlike *IBM1* loss-of-function mutants, *jmj26* did not show any morphological phenotype (Fig S8A), but mCHG levels increased significantly in RdDM-targeted transposons (Figs. 5B, C). Furthermore, *jmj26* showed increased mCHG in pericentromeric transposons, whereas *ibm1*-like hypermethylation was not observed in genic regions (Fig 5D, E). Gene expression was barely affected except for a few DNA methylation sensitive genes, including transposon genes (AT5G35935, AT4G01525) (Table S3). These observations support the functionality of *JMJ26* as a histone demethylase with different targets from *IBM1*, and also makes it a strong *a posteriori* candidate for causing the GWAS peak on the left end of chromosome 1.

### Major loci additively affect mCHG and influence transposon activity

Finally, we asked how the five major loci jointly shape mCHG variation at the population level. Although some of these genetic variants are clearly interacting at the molecular level (e.g., *MIR823A* directly regulates *CMT3*), the phenotypic effects appear to be additive (Fig. 6A), and jointly explain a large fraction of the variation: 30.5% and 14.8% of mCHG_|mCHH_ in RdDM-and CMT2-targeted transposons, respectively, while SNP-based heritabilities of these traits are 65% and 69%. mCHG levels decrease linearly with the number of mCHG-decreasing alleles, and the single line (out of 774) harboring mCHG-decreasing alleles at all five loci exhibits strong hypo-mCHG_|mCHH_ in RdDM-targeted transposons (Fig. 6A).

**Fig 6.**
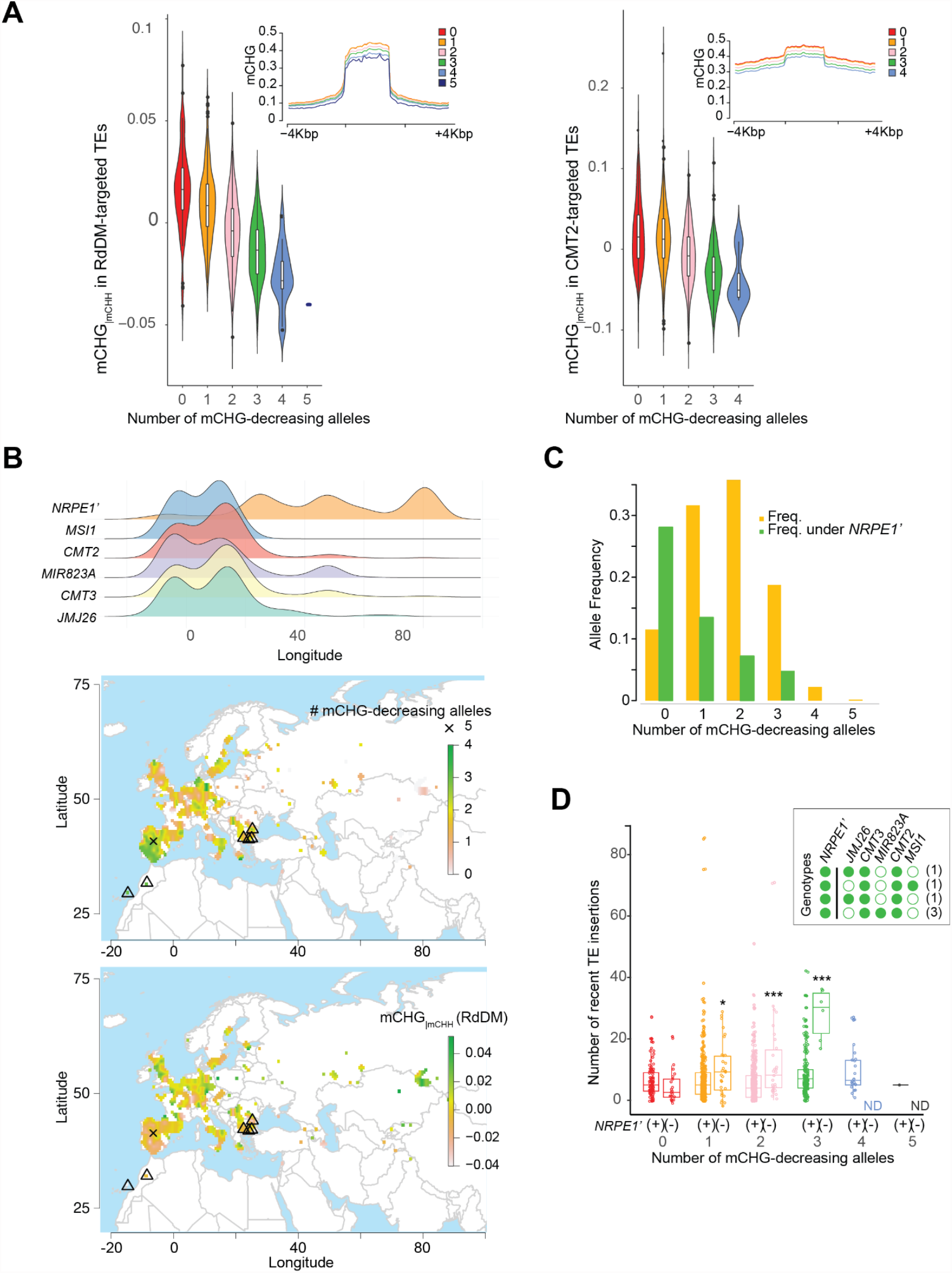
Cumulative effects and geographic distribution of mCHG-decreasing alleles. **(A)** Cumulative effect on mCHG_|mCHH_. Chr1:4035022 (*JMJ26*) was excluded from the analysis of CMT2-targeted transposons as it has no effect. Chr5:2355910 was used for *MSI1*. Inserted plots show mCHG levels around transposons, calculated using 20 sliding windows across the transposon body and flanking regions. **(B)** Longitudinal frequency distribution of mCHG-decreasing and *NRPE1’* alleles (top); geographical distribution of the number of mCHG-decreasing alleles (middle), and mCHG_|mCHH_ levels in RdDM-targeted transposons (bottom). Triangles correspond to lines in the inserted plot of panel D. **(C)** The frequency of distribution of the number of mCHG-decreasing alleles, separately for *NRPE1’* genotypes. **(D)** The number of recent transposon insertions (Baduel et al., 2021) as a function of *NRPE1’* genotype and the number of mCHG-decreasing alleles. Significance was tested using a permutation test (see methods; *** is *p* < 0.01, * is *p* < 0.1). The insert shows the six observed genotypes with three mCHG-decreasing and the *NRPE1’* non-reference allele. Filled circles indicate mCHH-or mCHG-decreasing alleles.

These five loci also affect the geographic distribution of mCHG for both RdDM-and CMT2-targeted transposons. In European lines (*n*=971), mCHG-decreasing alleles are almost exclusively found in the west, from 0 to 30ºE (Fig 6B, S9-10). Lines carrying more than three mCHG-decreasing alleles are rare (2.5 %), and the one line carrying all five such alleles is from the Iberian peninsula: IP-Mun-0 (9561), an admixed line of relict and western European descent (1001 Genomes Consortium, 2016).

Interestingly, the longitudinal gradient in the number of mCHG-decreasing alleles is opposite to the longitudinal cline of *NRPE1’* associated with lower mCHH (Sasaki et al., 2019) and higher transposon mobilization (Baduel et al., 2021) (Fig. 6B). No line was found carrying *NRPE1’* with more than three mCHG-decreasing alleles, and a permutation test shows that this negative correlation is unlikely to be due to population structure (Fig. 6C, S10C). A plausible alternative is some form of stabilizing selection on methylation levels: multiple mCHG-decreasing alleles in lines with lower mCHH levels might affect fitness analogously to double knockout mutants of the RdDM and CMT3 pathways that show pleiotropic morphological phenotypes and high transposon transcriptional activity (Cao and Jacobsen, 2002; Stroud et al., 2014; Chan et al., 2006).

To explore this further, we examined the effect of mCHG-negative alleles on transposon mobilization by intersecting our data with those of Baduel et al. (2021), who found a significant correlation between recent transposon mobilization and the *NRPE1’* non-reference allele. We found that this effect reflects an epistatic interaction with the five loci reported here: significantly increased transposon mobilization is only found in lines carrying the *NRPE1’* non-reference allele and two or more mCHG-decreasing alleles (*p*-value < 0.01; Figs 6D, S11). In particular, lines carrying three mCHG-decreasing alleles and the *NRPE1*’ non-reference allele have about thirty times more recent transposon insertions than lines carrying no mCHG-decreasing alleles and the *NRPE1’* non-reference allele. Such lines are not closely related and carry different transposon insertions, mostly Copia and MuDR. They are found in two distinct regions: north-western Africa, including the Canary Islands, and the south-eastern Balkans (Fig. 6B, Table S4).

## Discussion

### The genetics of non-CG transposon methylation

Non-CG methylation in plants is associated with gene silencing, especially for transposons. In previous studies, we identified an oligogenic architecture for mCHH on transposons involving major polymorphism at *CMT2, NRPE1, AGO1*, and *AGO9* — all involved in DNA methylation pathways (Kawakatsu et al., 2016; Dubin et al., 2015; Sasaki et al., 2019). Here we complement this with a conditional GWAS approach and identify five major polymorphisms affecting mCHG, presumably acting via methylation maintenance pathways: *CMT2* (different alleles), *CMT3, MIR823A* (directly regulating *CMT3*), *MSI1* (probably), and *JMJ26*. All had previously been shown to affect DNA methylation except *JMJ26*.

We observed major polymorphisms at several of the layers of regulation controlling mCHG, including methylation, de-methylation, and histone modification — even post-transcriptional regulation, in the form of a microRNA targeting the CMT3 pathway directly. The *DDM1* and *FAS1* alleles detected using non-stringent threshold would further contribute to the fine-tuning of DNA methylation over transposons genome-wide.

The genetic architecture revealed by these studies is very different from the “omnigenic” model (Boyle et al., 2017; Liu et al., 2019) which has come to represent a typical human GWAS result. It is also different from *trans*-regulation for reported human mCG variation (Hawe et al., 2022; Villicaña and Bell, 2021). Rather than a diffuse genetic architecture involving huge numbers of loci of small effect and unknown causal relationship to the phenotype, we find a small number of major loci with highly plausible mechanistic connection to the phenotype — and a diffuse background for which we lack power to dissect further. The results are reminiscent of those found for other clinal traits, like flowering time (Yeaman et al., 2016; Atwell et al., 2010), flower color (Ortiz-Barrientos, 2013), or eye and coat color in mammals (Miller et al., 2007; Beleza et al., 2013; Lloyd-Jones et al., 2017), suggesting that transposon methylation is likewise under selection, most likely as part of a plastic and environmentally sensitive genome defense system — a notion further supported by the association with transposon mobility reported here and be Baduel et al. (2021).

### The power and complexity of conditional GWAS

The performance of GWAS relies on using the right model for the relation between genotype and phenotype. As with other statistical methods, using the wrong model may lead to unpredictable results. However, lack of prior knowledge makes modeling difficult, and univariate linear models are therefore the default in GWAS. Here we focus on correlated traits. Correlations between traits may arise for a variety of reasons, mostly obviously a shared genetic basis (pleiotropy), and it is clear that such traits should, in principle, be analyzed jointly using some kind of multivariate analysis (Califano et al., 2012; Korte et al., 2012; Stegle et al., 2012; Stephens, 2013). Our results provide a dramatic illustration of the potential benefits of doing so. Despite high heritability, univariate GWAS of mCHG variation failed to detect any significant associations, leading us to conclude, erroneously, that the trait was simply too polygenic (Kawakatsu et al., 2016). In contrast, GWAS of the same data using a conditional model that controlled for mCHH revealed an oligogenic architecture with five major loci — qualitatively similar to what we had previously seen for mCHH using univariate GWAS, also in that it mainly identified loci corresponding to biologically meaningful *a priori* candidates (Kawakatsu et al., 2016; Dubin et al., 2015; Sasaki et al., 2019).

However, while our conditional GWAS approach clearly improved power to reject the null hypothesis of no association, interpreting the result in terms of causality is more difficult. We believe that, by controlling for mCHH, we have effectively simplified the trait, revealing genetic factors affecting mCHG only, perhaps by affecting the maintenance of this type of DNA methylation. There is considerable background experimental evidence to support such a model. For example, the RdDM pathway would affect both mCHH and mCHG via *de novo* methylation, while mCHG (but not mCHH) would be also maintained by the CMT3 pathway, and by controlling the former, we would reveal the latter.

Our GWAS results are consistent with this interpretation. Polymorphism at *NRPE1*, a key component of the RdDM pathway, was revealed in univariate GWAS, while variation in *CMT3* (and its regulator, *MIR823a*) was only found after controlling for mCHH. At the same time, it seems clear that reality is more complex, and that both mCHH and mCHG are regulated by multiple homeostatic mechanisms that also involve factors not included in our model, like histone modifications (Zhang et al., 2021). It is therefore not surprising that most of the polymorphisms we have identified seem to affect both traits, albeit not to the same extent (exceptions include the one at *MIR823A*, which only seems to affect mCHG, and the previously identified *CMT2a’* and *CMT2b’* polymorphisms, which only affect mCHH on CMT2-targeted transposons; see Figs S2, S3).

Two other GWAS models produced consistent results, but had less power (Fig S12, S13). A fully parameterized multi-trait mixed model for mCHH and mCHG (MTMM; see Korte et al., 2012) confirmed that the new associations presented in this paper were “specific” to one of the two traits, while a conditional model for mCHH while controlling for mCHG (mCHH_|mCHG_) also identified most of these loci, plus the previously identified *CMT2* associations (Dubin et al., 2015; Sasaki et al., 2019), which affect mCHH only.

In conclusion, we agree with Stephens (2013) that multi-trait association methods can have much greater power than univariate methods, but require an appropriate statistical framework for interpretation. However, in a model organism like *A. thaliana*, further experiments should guide the construction of such a framework.

### Functional importance of non-CG transposon methylation

While genomics has revealed striking geographic variation in DNA methylation, and an equally striking genetic architecture underlying this, the functional importance of all this variation has been rather less clear. Based on decades of molecular biology studies, it is reasonable to speculate that it must play some role in regulating transposon activity (Stroud et al., 2013; Kim and Zilberman, 2014; Pikaard and Mittelsten Scheid, 2014; Matzke et al., 2015). Non-CG methylation is part of a redundant system with strong epistasis between mCHH and mCHG in keeping transposons transcriptionally silent (Cao and Jacobsen, 2002; Stroud et al., 2014; Chan et al., 2006). However, direct evidence based on transposon mobility in nature has been missing. Now, the evidence is gathering that the allelic variants responsible for transposon methylation variation do indeed affect transposon mobility. Baduel et al (2021) showed that the *NRPE1’* polymorphism identified as affecting mCHH on RdDM-targeted transposons by Sasaki et al (2019) is associated with recent transposon mobility, and we show here that the multiple loci controlling mCHG appear to play a similar role. Our findings connect complex genome defense systems proposed in molecular biology studies with natural populations. Furthermore, the geographic distribution of the relevant multi-locus genotypes suggests that selection may be acting to maintain the appropriate level of transposon silencing — for unknown reasons. Further studies, such as crosses with new allele combinations and common garden experiments, will be required to address these questions.

## Materials and methods

### Analyzed data sets

#### DNA methylation data

The DNA methylation data sets are summarized in Table S5. Briefly, we analyzed a bisulfite-sequencing data set published in the 1001 epigenome project (Kawakatsu et al., 2016), 85 epigenetic-related mutants (Stroud et al., 2013), and our own sequence data described below. All reads were mapped on the appropriate pseudo-genome provided by the 1001 genome project (1001 Genomes Consortium., 2016) using a Methylpy pipeline v1.2 (Schultz et al., 2015). DNA methylation levels were estimated as weighted methylation levels for each transposon defined in Araport11 annotation. CMT2-and RdDM-targeted transposons were defined as having differential levels of methylation (>0.1) between wild-type and *cmt2* or *drm1drm2* in Col-0 previously described in Kawakatsu et al. (2016). For each line, average DNA methylation was calculated using all transposons for which at least one read was mapped.

#### RNA-seq data

The RNA-seq data are summarized in Table S5. Quality control and adapter trimming of all RNA-seq data were conducted using FASTP (Chen et al., 2018). Adapter trimmed reads were aligned on the reference genome (Araport11; Cheng et al., 2017) using STAR v2.7 (Dobin et al., 2013) with default settings. For all annotated genes, mapped read counts were calculated using featureCounts v2.0.1 in the Subread package (Liao et al., 2013), accepting reads that were mapped on multiple genes (option -M). Using the count data, differentially expressed genes were detected by edgeR v3.1 (Robinson et al., 2010) with glmQLFTest() function (FDR < 0.05). For transposon expression, reads were mapped as described above but allowing multiple hits (STAR --outFilterMultimapNmax 100 and --winAnchorMultimapNmax 100). Mapped reads were calculated by TEtoolkits (Jin et al., 2015) using the TEcount() function, and differentially expressed transposons were detected using edgeR. *CMT3* expression in natural populations were downloaded from the 1001 epigenome project data (Kawakatsu et al., 2016).

### Statistics

#### Genome-wide association studies

Univariate and conditional GWAS were carried out using LIMIX (Lippert et al., 2014) version

3.0.4 with a full genome SNP matrix for 774 lines (Kawakatsu et al., 2016) from the 1001 genome project (10,709,949 SNPs) and the following linear mixed model (LMM),

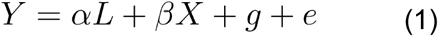

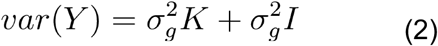

where *Y* is the *n* × *1* vector of a phenotype, fixed terms, *L* and *X*, are the *n* × *1* vectors corresponding to a covariate and a genotype to be tested (SNP) with the parameters *α* and *β*, respectively. 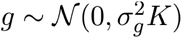 and 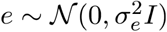 are random terms, including the identity-by-state kinship matrix *K* representing the genetic relatedness (Yu et al., 2006; Kang et al., 2008) and the residuals, respectively. *Y, L, X* and were z-scored. Univariate models without cofactor take *α=0*. SNPs that satisfied minor allele frequency > 5% were used for association studies. Bonferroni-correction was used for multiple-testing correction after excluding all SNPs less than MAF 5%. Multi-trait linear mixed models (Fig. S12) were performed to identify common genetic effects (common) and differing genetic effects (specific) between two traits using LIMIX (Lippert et al., 2014) following models described in Korte et al. (2012).

#### Enrichment test

To assess GWAS results, we calculated FDR and enrichment of *a priori* gene for each GWAS result (as described in Atwell et al. (2010)) using a list of 79 epigenetic regulators from Kawakatsu et al. (2016). The most significant *p*-value within 15 kb of a gene (MAF > 5%) was assigned as the significance of the gene.

#### Heritability

SNP-heritability was estimated using REML implemented in mixmogam (https://github.com/bvilhjal/mixmogam/blob/master/phenotypeData.py). The proportion of phenotypic variation explained by identified genetic variants was calculated using *r*^2^ (Nakagawa and Schielzeth, 2013) using R scripts published in Sasaki et al. (2018).

#### Correlation of the allelic effects and molecular phenotypes

For 9329 transposons common across 774 lines, differential mCHG levels induced by alleles were estimated for each transposon as the differential average methylation level between lines carrying the reference (Col-0) and the alternative allele. Differential mCHG levels induced by mutants were calculated analogously (Stroud et al., 2013). The details are described in Sasaki et al. (2019).

#### Gene effect of loss-of-function mutants

Gene functions of *MIR823A* and *JMJ26* were tested using transgenic lines described below. We measured DNA methylation levels for two or three independently grown plants, and the significance of differential mean values across RdDM-and CMT2-targeted transposons was tested by Welch’s t-test (two-tailed). DNA methylation pattern of *jmj26* was compared to *ibm1* and the control samples from published data sets (Stroud et al., 2013).

#### Permutation tests

We evaluated effects of mCHG-decreasing alleles using permutation tests with 3000 randomly chosen SNP-sets with the same allele frequencies as the identified alleles along the genome. For correlations between number of mCHG-decreasing alleles and longitude or linkage disequilibrium between the five alleles and *NRPE1’* alleles, we compared the observed to the 3000 randomly chosen samples directly. For evaluation of epistatic effects between *NRPE1’* and mCHG-decreasing alleles on transposon activity, we randomly chose 3000 SNPs with the same allele frequency as *NRPE1’* and tested the effect of zero to five mCHG-decreasing alleles.

### Plant materials

Plants were grown at the 21°C with a 16-h light/8-h dark cycle and humidity 60%. Whole plant rosettes were harvested individually when they reached the 9-true-leaf stage of development. The t-DNA insertion line of *jmj26* (SALKseq_069498.1) and a gamma-ray mutant *msi1-2* (Jullien et al., 2008) were purchased from Nottingham Arabidopsis Stock Center (http://arabidopsis.info/) for the functional analysis. Homozygous *jmj26* was used for bisulfite sequencing after the second generation. *msi1-2* was maintained as a segregating population, and heterozygous lines were selected by genotyping and measuring *MIS1* expression by qRT-PCR.

### Transgenic lines

#### CRISPR/CAS9 mutant lines

Loss-of-function mutants *mir823a* for Col-0 and 9771 were generated using CRISPR/CAS9 methods described in Xing et al (2014). Briefly, to generate a vector carrying two sgRNAs targeting *MIR823A*, the target sites were incorporated into forward and reverse PCR primers, and the fragment amplified from pCBC-DT1T2 (Table S6). Subsequently, an amplified and purified fragment was assembled using the Golden Gate reaction into pHSE401E modified to contain a mCherry marker. The resulting vector was transformed into *Agrobacterium tumefaciens* GV3101/pSOUP, then transformed into *A. thaliana* by the floral dip method (Zhang et al., 2006). T1 seeds were screened by mCherry marker under the fluorescence stereomicroscope (Discovery.V8, Zeiss), and the mutations were genotyped by Sanger sequencing. T2 seeds without mCherry signal were kept to amplify the stable T3 generations, which were used for further analyses.

#### Rescue analyses

The gene function of *JMJ26* was confirmed by rescue experiments. The protein-coding genomic sequence including 3 kbp promoter sequence was PCR-amplified and cloned into the pGreen0029 binary vector (Hellens et al., 2000) using In-Fusion Cloning system (Takara Bio Europe, Saint-Germain-en-Laye, France) for Col-0 and DraIV 6-22 (5984) carrying the reference and the alternative allele, respectively. These constructs were used for floral dipping transformation to Col-0 background *jmj26* (SALKseq_069498.1) to create transgenic lines (Zhang et al., 2006).

### mRNA abundance

Total RNA was extracted using ZR Plant MiniPrep Kit (Zymo Research) including a DNase I treatment and was quantified by fluorometer Qubit 4 (Invitrogen). cDNA was synthesized using the SuperScript III First-Strand Synthesis System (Invitrogen) according to the manufacturer’s protocols. qRT-PCR was performed using the LightCycler 96 system (Roche) with FastStart Essential DNA Green Master (Roche). The transcript abundance was estimated by ddCT methods with an internal control *SAND* (AT2G28390) (Czechowski et al., 2005). Primer information is listed in Table S6.

### MicroRNA abundance

#### Material collection and RNA extraction

Siliques were collected five days after pollination when the embryos reached an early torpedo stage according to the previous report (Papareddy et al., 2021). Using the tungsten needles, seeds were dissected from siliques and collected in 1.5 ml tubes, and washed 3-4 times with fresh 10% RNA later solution (Thermo Fisher Scientific). After the last wash, the buffer was replaced with 500 µl TRIzol™ Reagent (Ambion) and total RNA was isolated as described in Hofmann et al. (2019).

#### qRT-PCR Analysis

The quantification was conducted according to the protocol in Papareddy et al. (2021). Briefly. cDNA synthesis was performed using SuperScript III System (Invitrogen), and the corresponding stem-loop primers were added to the reverse transcription reaction. qRT-PCR analysis was performed on the LightCycler 96 Instrument (Roche) with the FastStart Essential DNA Green Master (Roche). The transcript abundance was estimated by ddCT methods with an internal control *miRNA160*.

### RNA sequencing

RNA was extracted in the same protocol with qRT-PCR for mRNA.Total RNA was treated with the poly(A) RNA Selection Kit (Lexogen). Libraries were prepared according to the manufacturer’s protocol in NEBNext Ultra II Directional RNA Library Prep Kit (New England BioLabs) and individually indexed with NEBNext Multiplex Oligos for Illumina (New England BioLabs). The quantity and quality of each amplified library were analyzed by using Fragment Analyzer (Agilent) and HS NGS Fragment Kit (Agilent).

### Bisulfite sequencing

Genomic DNA was extracted using GeneJET Plant Genomic DNA Purification Kit (Thermo Scientific) and sheared by E220 Focused-ultrasonicator (Covaris) to target average fragments around 350 bp. Sequencing libraries were prepared using NebNext Ultra II DNA Library Prep Kit (New England BioLabs) with Methylated Adaptor (New England BioLabs). Adapter-ligated DNA was carried through the bisulfite conversion using EZ-96 DNA Methylation-Gold™ MagPrep Kit (Zymo Research). Bisulfite-treated samples were amplified by EpiMark Hot Start Taq DNA Polymerase and indexed with NEBNext Multiplex Oligos for Illumina (New England BioLabs). All libraries were sequenced by Illumina NextSeq550 or Hiseq2500.

## Data availability

All data are summarized in Table S5. Sequencing data have been deposited to GSE194406.

## Acknowledgments

We are grateful to Fred Berger, Tetsuji Kakutani, Arturo Marí-Ordóñez, Ortrun Mittelsten Scheid and all member of the Nordborg lab for discussions, and to Pierre Baduel, Vincent Colot, and Leandro Quadrana for discussions and sharing of transposon mobilization data. We thank Pierre Baduel, Pieter Clauw, Vincent Colot, Thomas Ellis, and Bob Schmitz for extremely helpful comments on the manuscript, Jian-Kang Zhu and Kai Tang for sharing *ros3* seeds and methyl-C seq data, and Rahul Pisupati for help with data analyses. The bisulfite and RNA sequencing were performed by the Next Generation Sequencing Facility at Vienna BioCenter Core Facilities (VBCF), a member of the Vienna BioCenter (VBC), Austria. This work was funded in part by ERC AdvG 789037 EPICLINES to MN.

## Supplemental tables

Table S1. Linkage disequilibrium between five loci

Table S2. *a priori* genes identified by enrichment tests (FDR=0.2) Table S3. The list of genes significantly affected in *jmj26*

Table S4. Lines carrying three mCHG-decreasing alleles with mCHH-decreasing *NRPE1’* allele

Table S5. Key resources

Table S6. Primer list

## Supplemental figures

**Fig S1.**
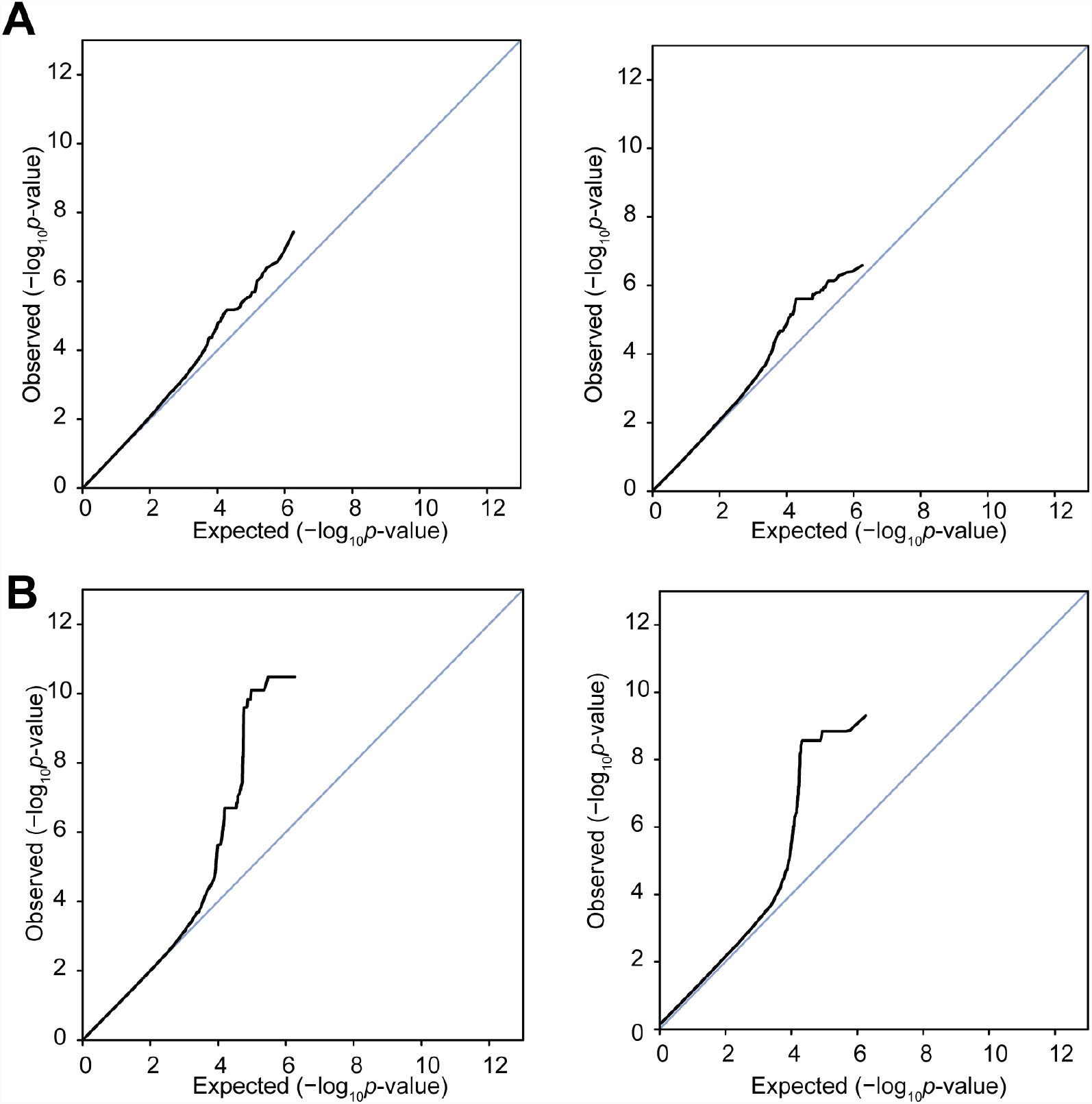
Distribution of p-values for two GWAS models. QQ plots for univariate models for mCHG levels **(A)** and conditional models for mCHG_|mCHH_ **(B)** in RdDM-targeted transposons (left) and CMT2-targeted transposons (right).

**Fig S2.**
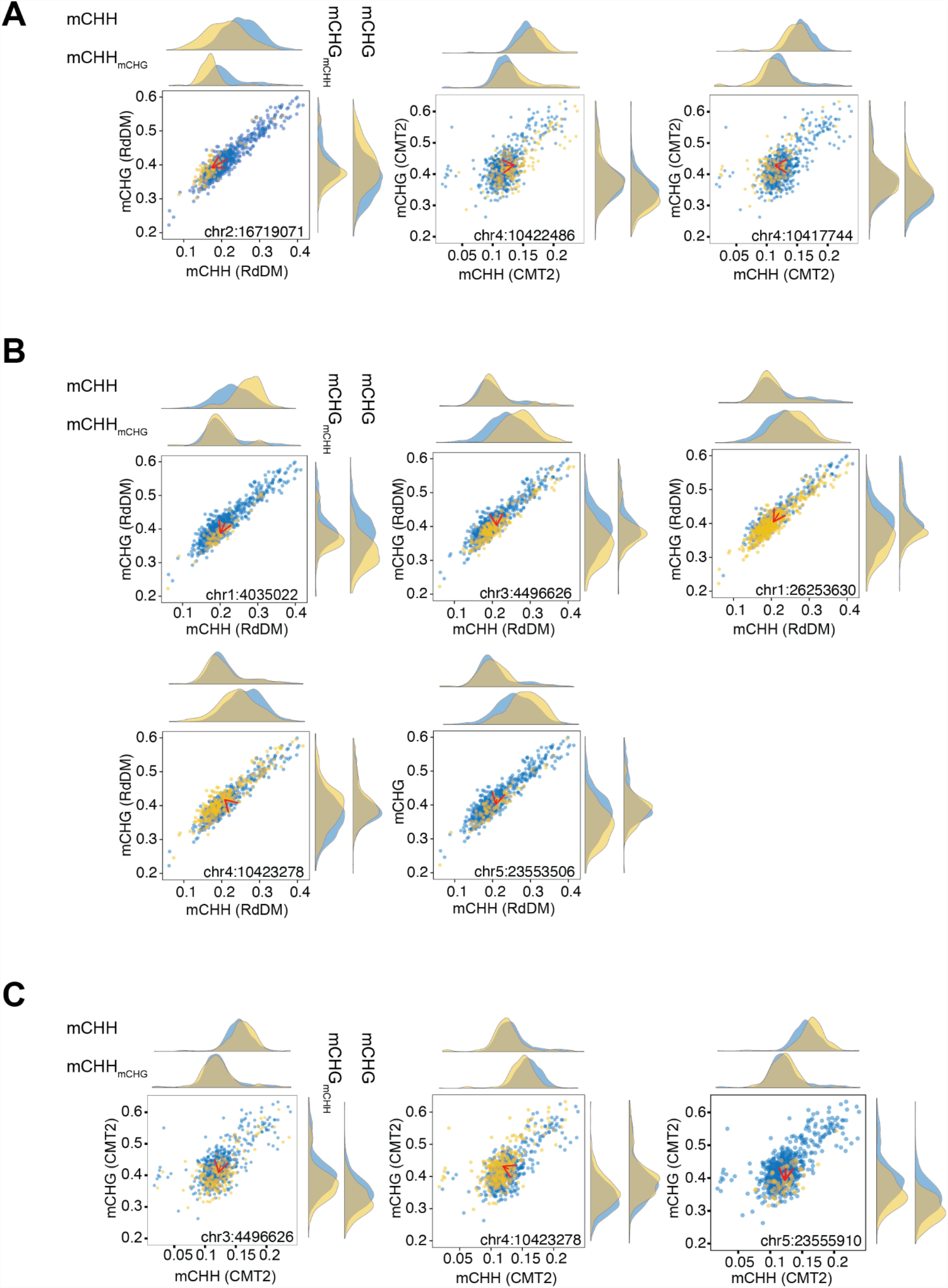
Distribution of phenotypes and the allelic effects. The scatter plots illustrate the allelic effects on mCHH (**A** from Sasaki et al., 2019) and mCHG|mCHH in RdDM-**(B)** and CMT2-targeted transposons **(C)**. Marginal phenotypic distributions for the reference and the alternative allele are plotted in blue and yellow, for univariate and conditional distributions. Red vectors show shifts of the mean value from the reference to the alternative allele in 2-dimensional phenotype space.

**Fig S3.**
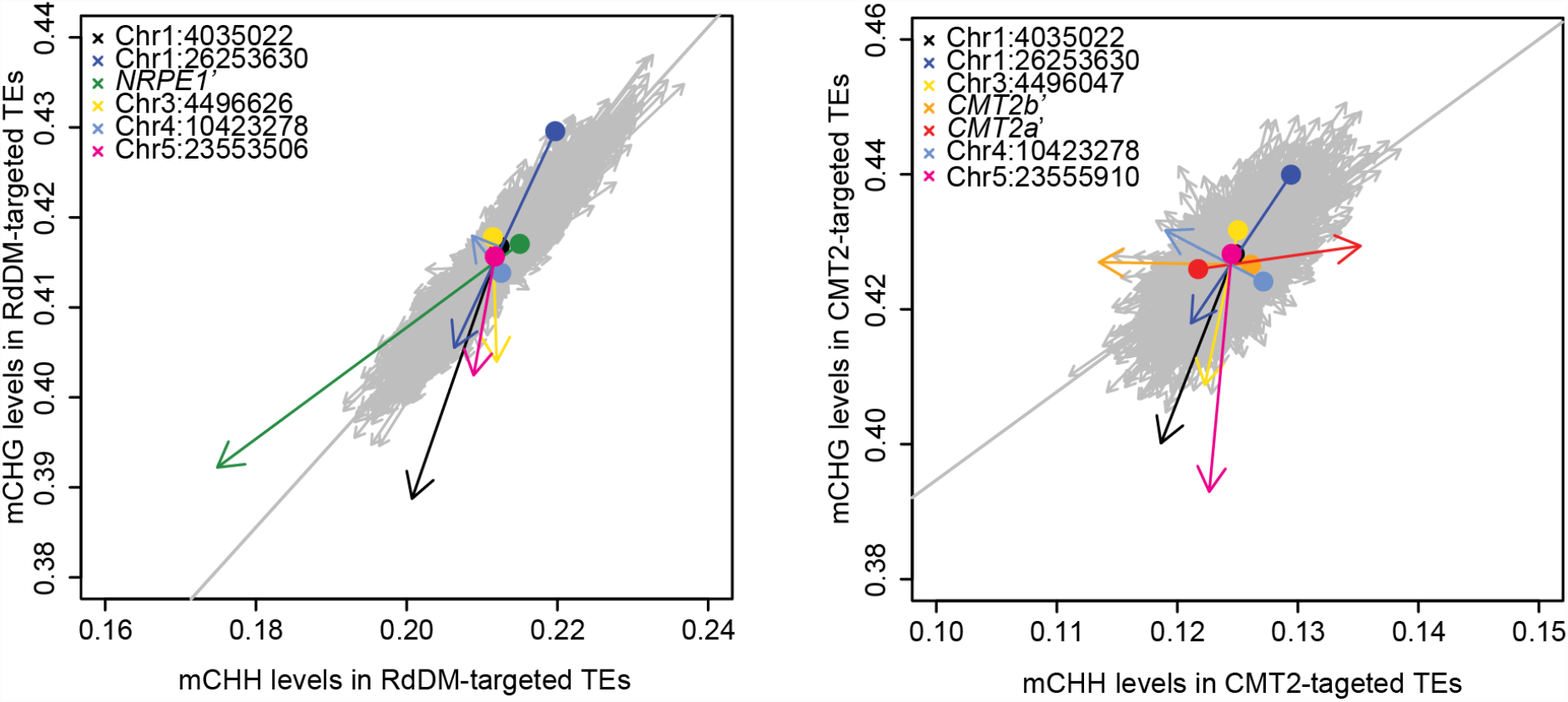
Comparison of the allelic effects on mCHH and mCHG. Each arrow shows the mean value shift of mCHH and mCHG levels from reference to the alternative allele. Left and right plot correspond to RdDM-and CMT2-targeted transposons, respectively. Gray arrows correspond to the genetic effects of randomly sampled SNPs, keeping the allele frequency of each tested SNP (x 3000 for each allele). Diagonal lines correspond to linear regression lines of mCHH and mCHG.

**Fig S4.**
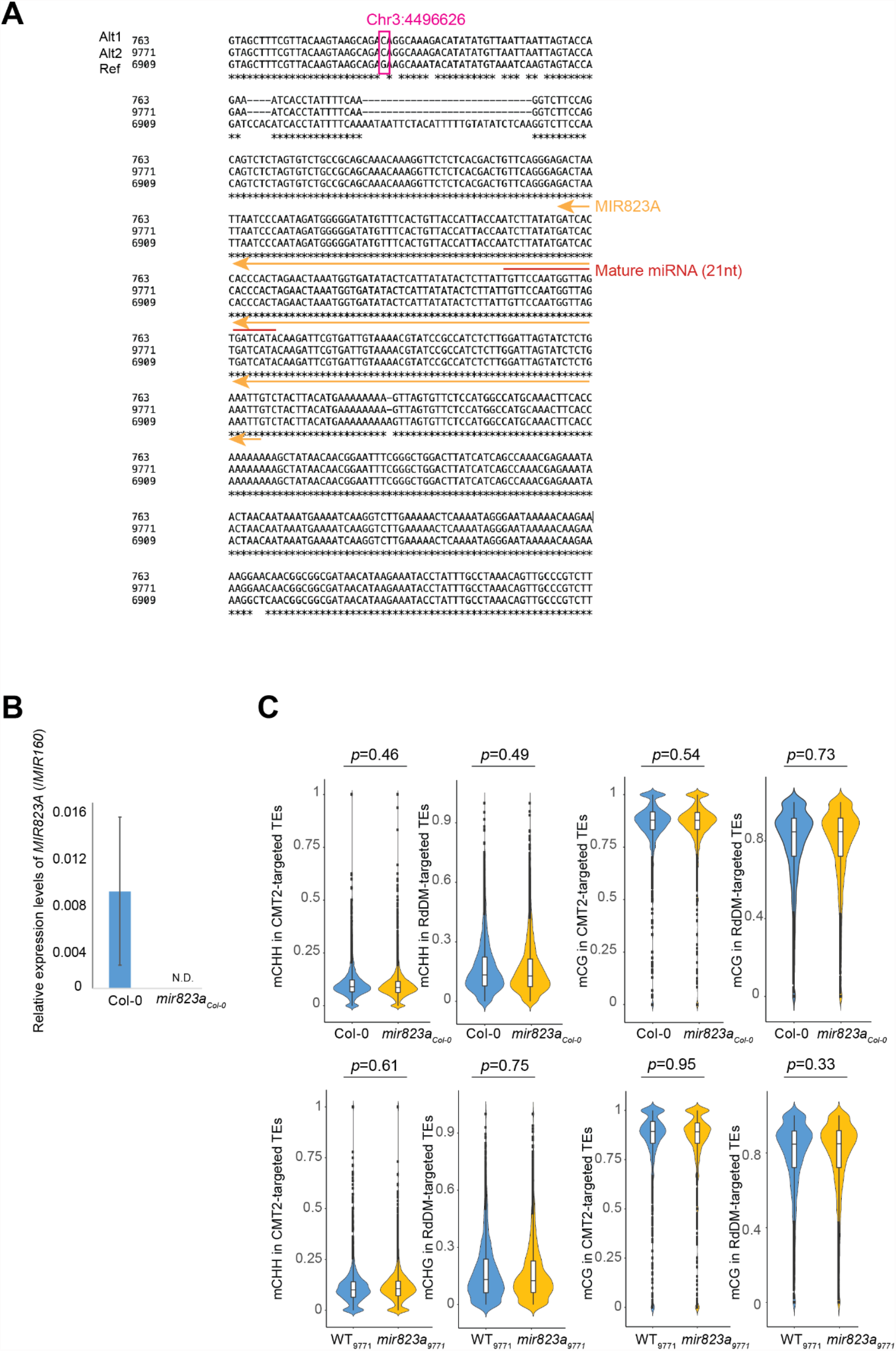
Characterization of *MIR823A*. **(A)** Sanger sequences around *MIR823A* region for lines carrying reference allele (Col-0) and the alternative allele (763, 9771). **(B)** *MIR823A* expression in a CRISPR/CAS9 *mir823a* mutant (Col-0) by qRT-PCR. Error bar shows standard deviation (*n*=3). **(C)** Effects of *MIR823A* on mCHH and mCG levels in RdDM and CMT2-targeted transposons in *mir823a* Col-0 and 9771 background.

**Fig S5.**
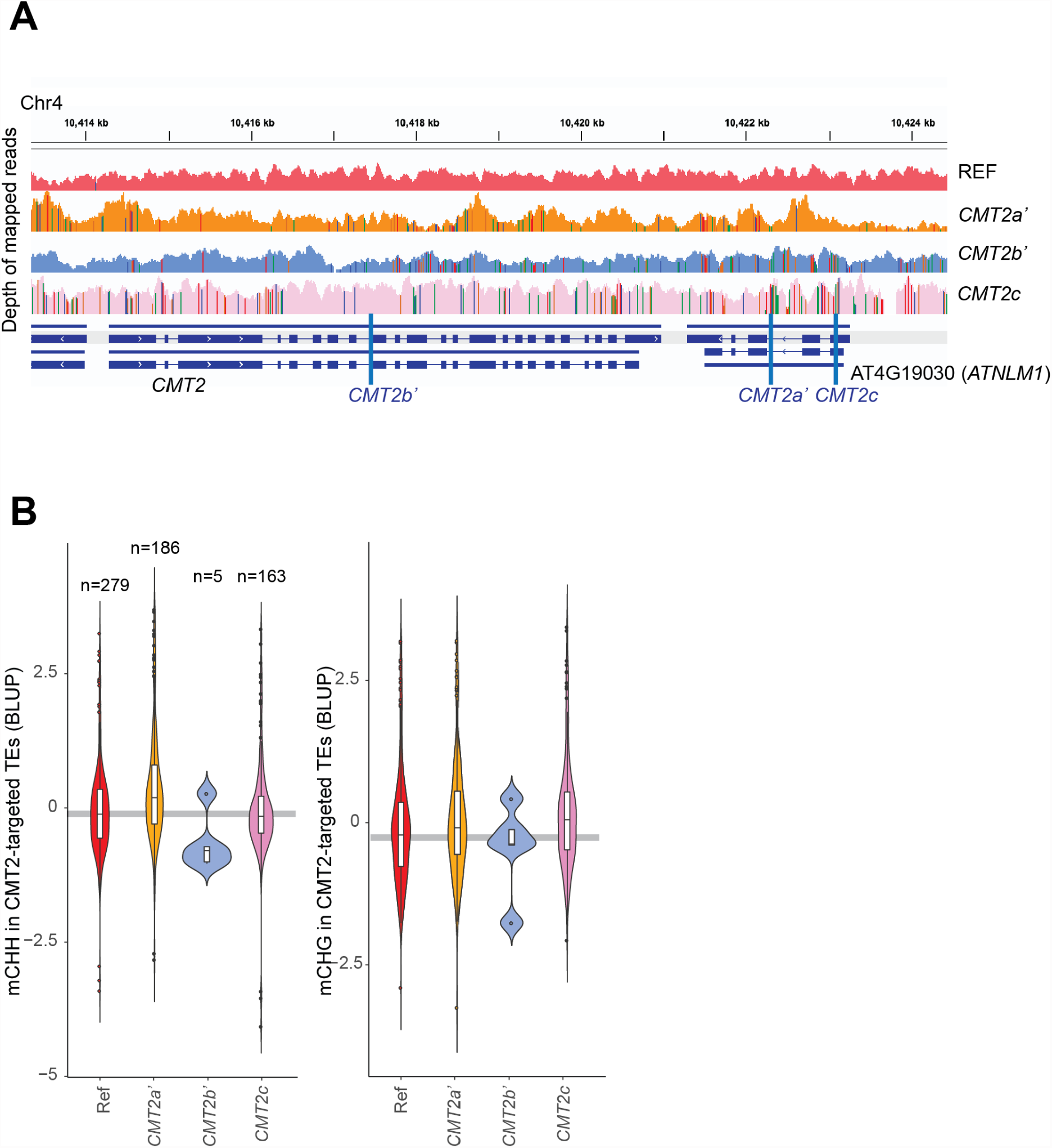
The effects of *CMT2* alleles on non-CG methylation. **(A)** Genome structure of three CMT2 alleles associated with non-CG methylation variation. The CMT2 region was illustrated by mapped short-read DNA-seq data (IGV browser) for reference line (Col-0), *CMT2a’* (10018), *CMT2b’* (6969), and *CMT2c* (10023). Vertical colored lines in the IGV plots indicate SNPs. **(B)** The allelic effects on genome-wide average mCHH and mCHG levels in CMT2-targeted transposons. Only lines carrying one allele were compared with the reference line. Horizontal gray lines show median values of the reference lines.

**Fig S6.**
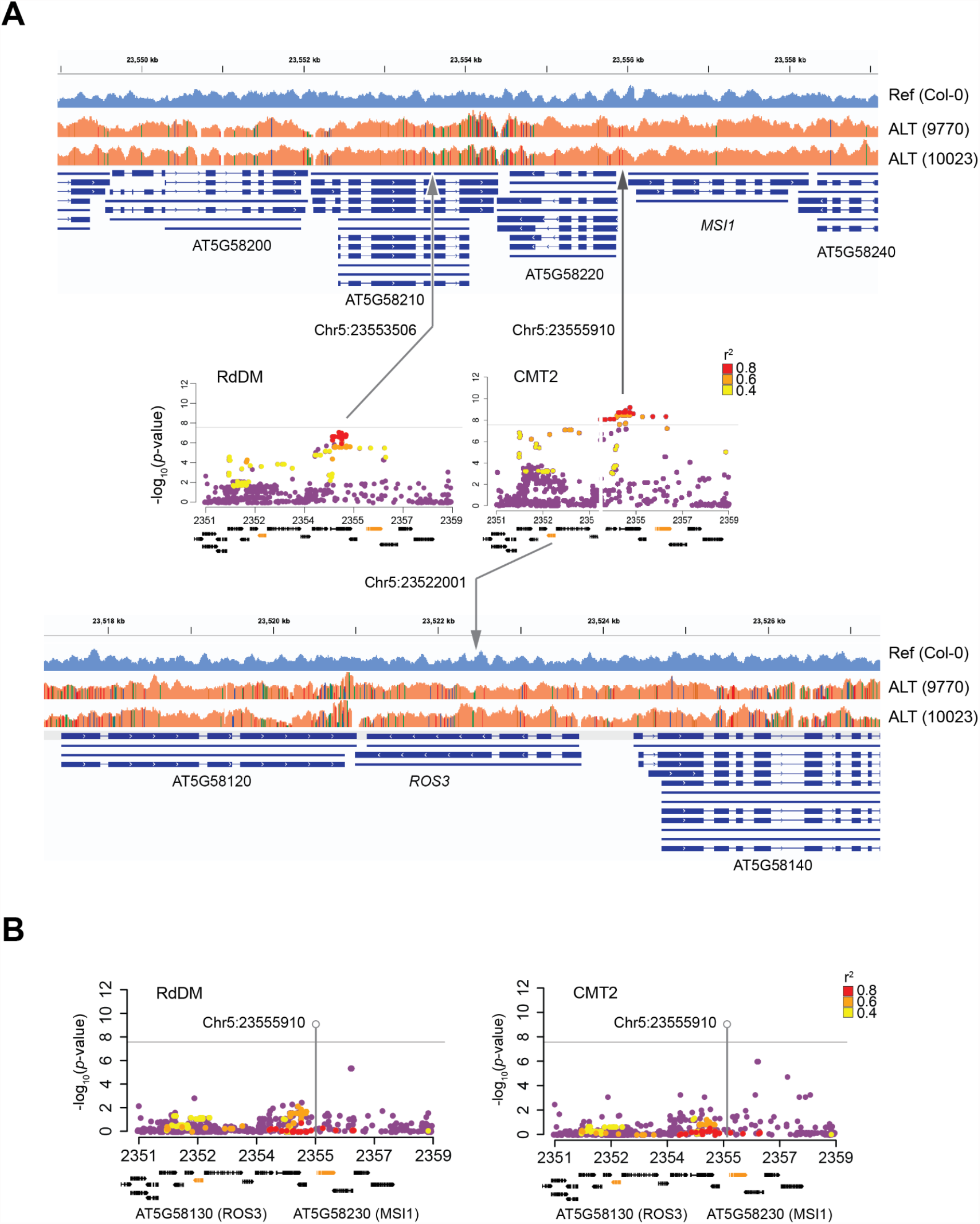
Genetic variation around *MSI1 and ROS3*. **(A)** Zoom-in Manhattan plots (Fig 3) and the genome structure around Chr5:23553506, 23555910 (top), and 23522001 (bottom) illustrated by mapped short-read DNA-seq data (IGV browser). Vertical colored lines in the IGV plots show SNPs. **(B)** Conditional GWAS for mCHG in RdDM-and CMT2-targeted transposons. mCHH and Chr5:23555910 were both used as co-factors. Gray vertical lines indicate the Chr5:2355910 position, and horizontal lines show the genome-wide significance (p=0.05 by Bonferroni correction). *r*^2^ was calculated from chr5:23553506 and chr5:23555910 for mCHG_RdDM_ and mCHG_CMT2_, respectively.

**Fig S7.**
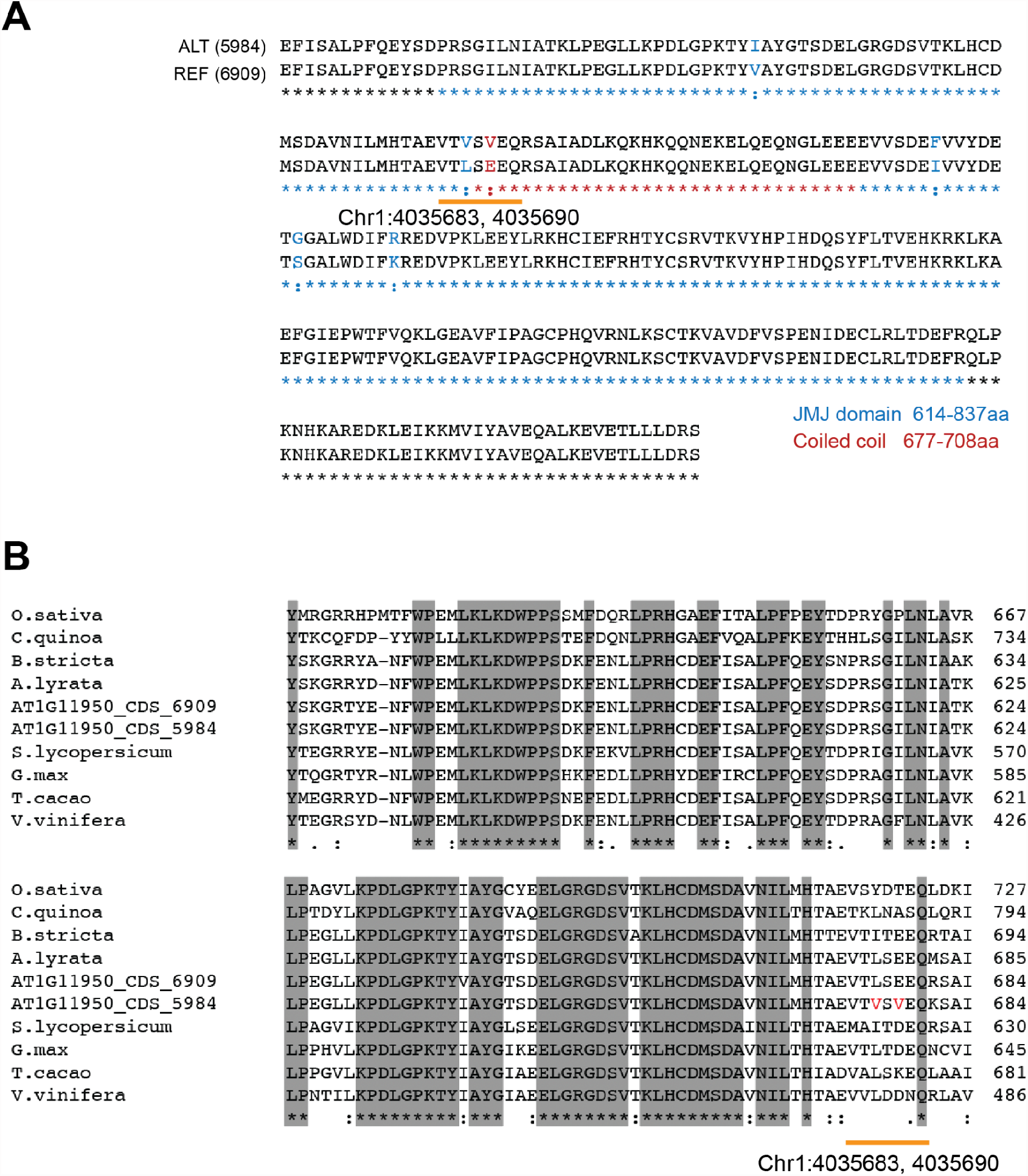
Characterization of *JMJ26* allele. Predicted amino acid sequences around jmjC domain of reference (6909) and the alternative allele (DraIV 6-22)(**A**) and the conserved region (**B**). Orange bars indicate nonsynonymous mutations associated with mCHG_|mCHH_ in RdDM-targeted TEs. The sequences were predicted based on polymorphism data provided by the 1001 genome project. The domain information follows the TAIR10 annotation.

**Fig S8.**
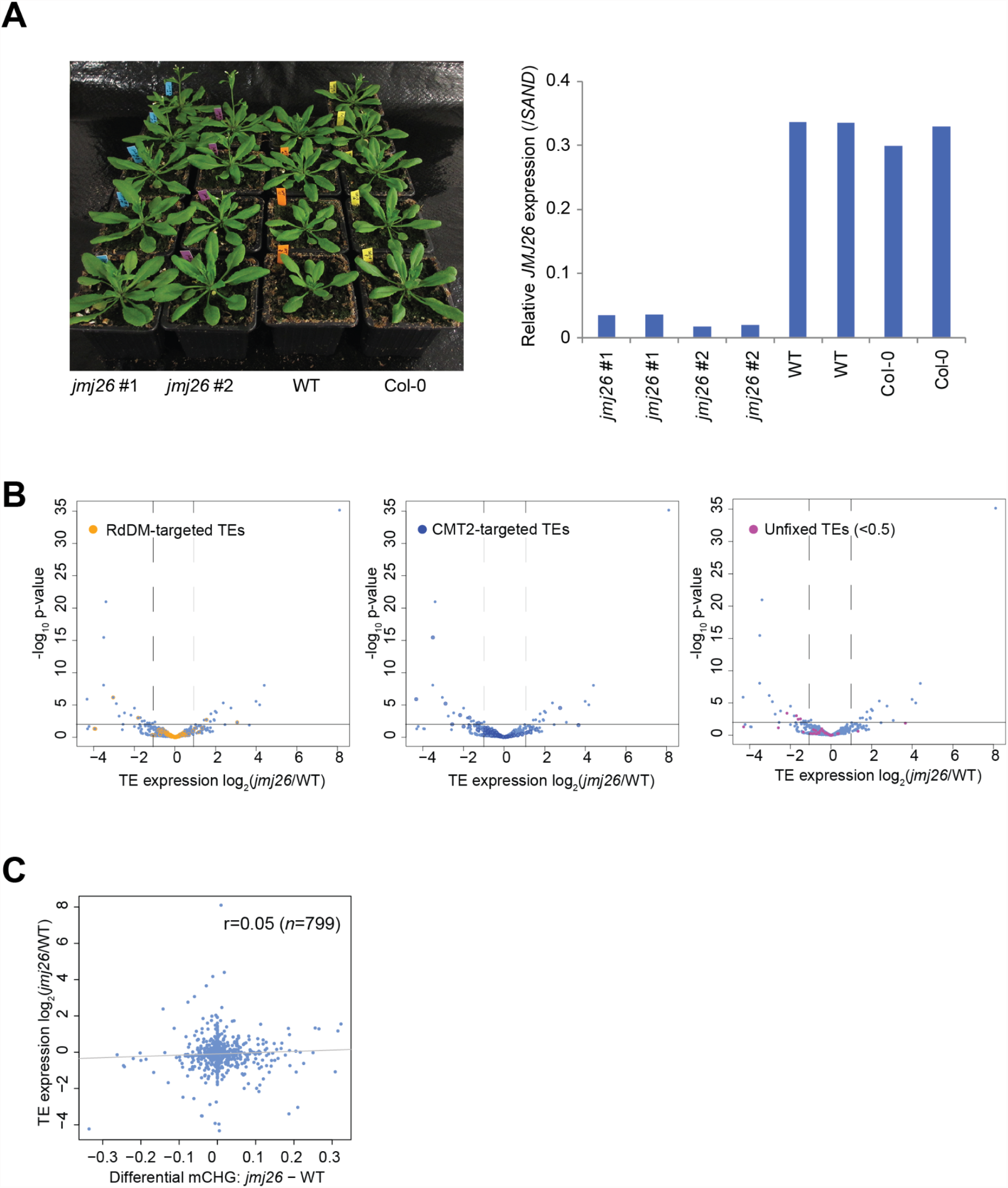
Molecular phenotypes of *jmj26*. **(A)** Characterization of loss-of-function mutants, *jmj26*. Both #1 and #2 were *jmj26* homozygous lines isolated from SALKseq_069498.1 and propagated separately. WT is a segregated line carrying active *JMJ26* in the same stock. Morphology of *jmj26* and Col-0 (left) and *JMJ26* expression in leaves of *jmj26* and the wild type (right). Expression was measured by qRT-PCR. **(B)** Volcano plots show the effects on transposon transcripts highlighted RdDM-and CMT2-targeted transposons. **(C)** The scatter plot shows the effect of mCHG methylation on transposon transcripts. The gray line shows the linear regression line.

**Fig S9.**
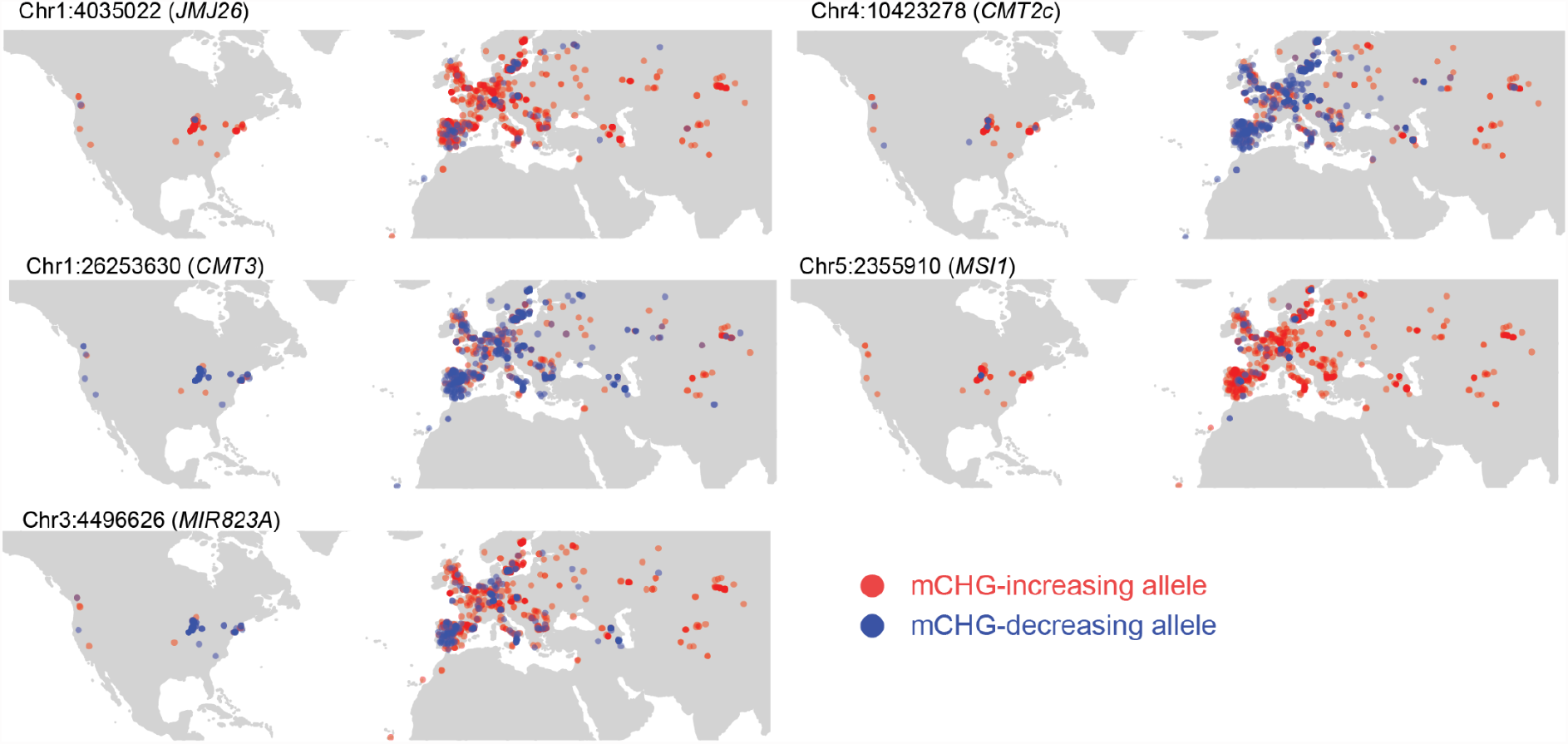
Geographical distribution of mCHG-decreasing alleles. The plots show the origin of lines carrying mCHG-increasing or decreasing alleles.

**Fig S10.**
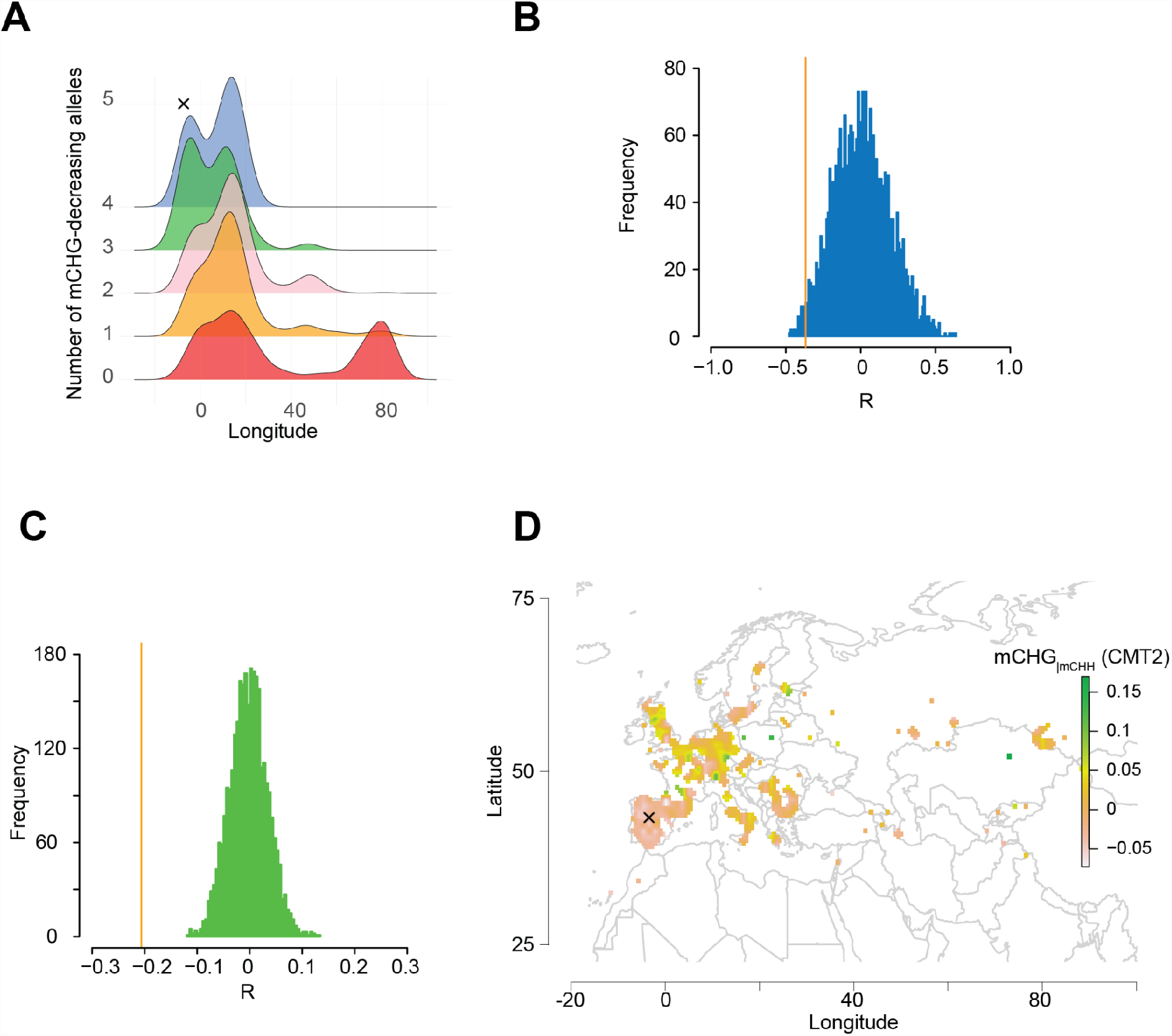
Geographical distribution of cumulative mCHG-decreasing alleles. **(A)** Longitudinal frequencies of cumulative mCHG-decreasing alleles (corresponding to five alleles in Fig. 6A). **(B)** The histogram shows how the cumulative mCHG-decreasing allele number is correlated with longitude of the origins. The blue histogram shows permuted Pearson’s correlation coefficients (R) between numbers of mCHG-decreasing alleles and longitude of the origin, maintaining the allele frequencies. The permutation tests were repeated 3000 times for 971 lines ranging from longitude -20º to 100º. The orange vertical line indicates the observed value. **(C)** Histogram similarly shows permuted Pearson’s correlation coefficients between number of mCHG-decreasing alleles and *NRPE1’* genotype. The orange line is the observation. mCHH-increasing and decreasing *NRPE1’* alleles are 0 and 1, respectively. **(D)** The geographic distribution of mCHG_|mCHH_ levels in CMT2-targeted transposons.

**Fig 11.**
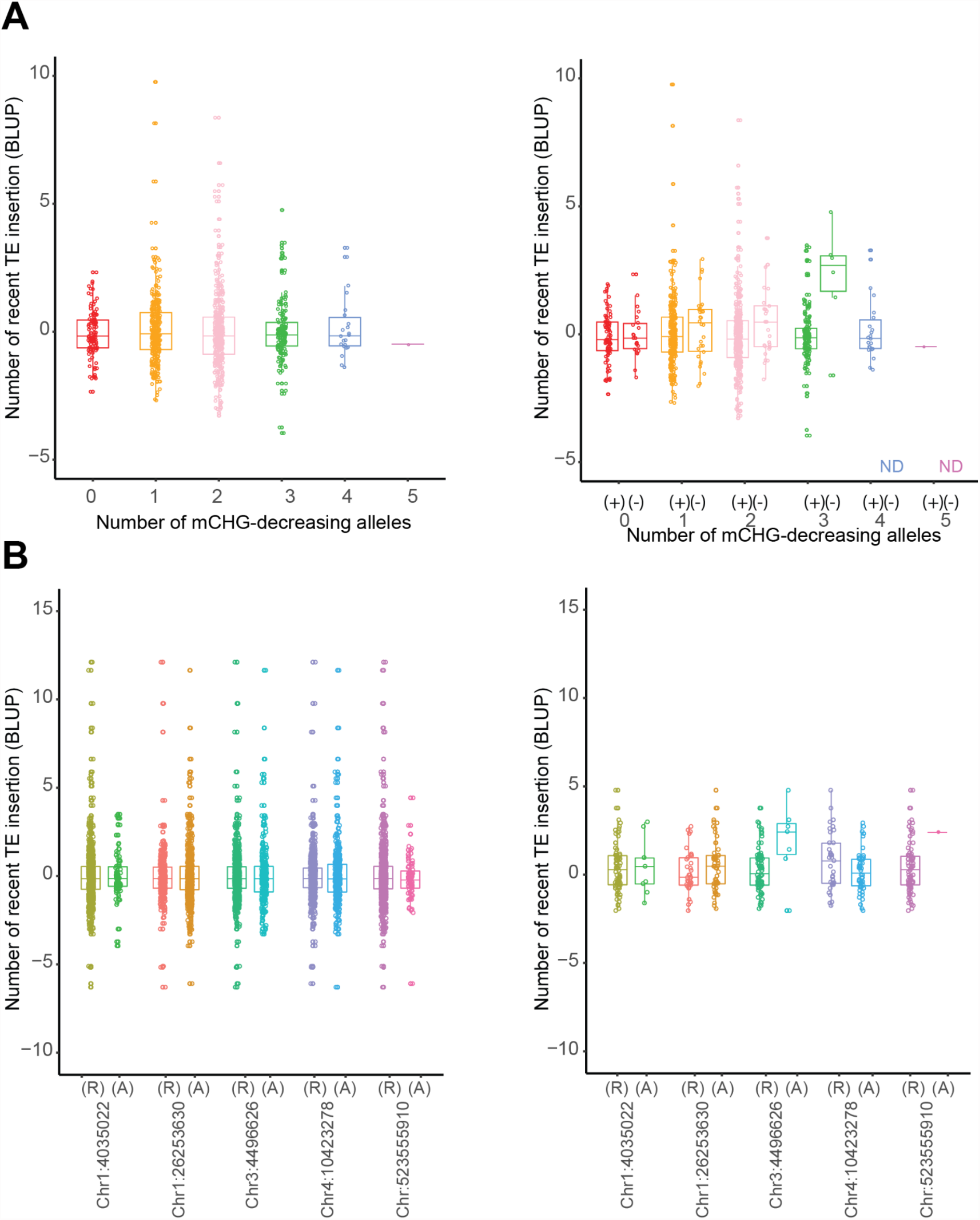
Function of mCHG-decreasing alleles on transposon activities. **(A)** Effects of the cumulative mCHG-decreasing alleles in whole populations (left) and combination with *NRPE1’* allele (right). (+) and (-) are lines carrying *NRPE1’* reference (mCHH-increasing) and the alternative (mCHH-decreasing) alleles, respectively. **(B)** Effects of the five major mCHG-decreasing alleles with *NRPE1’* reference (left) and the alternative (right) populations. (R) and (A) are reference and the alternative alleles, respectively.

**Fig S12.**
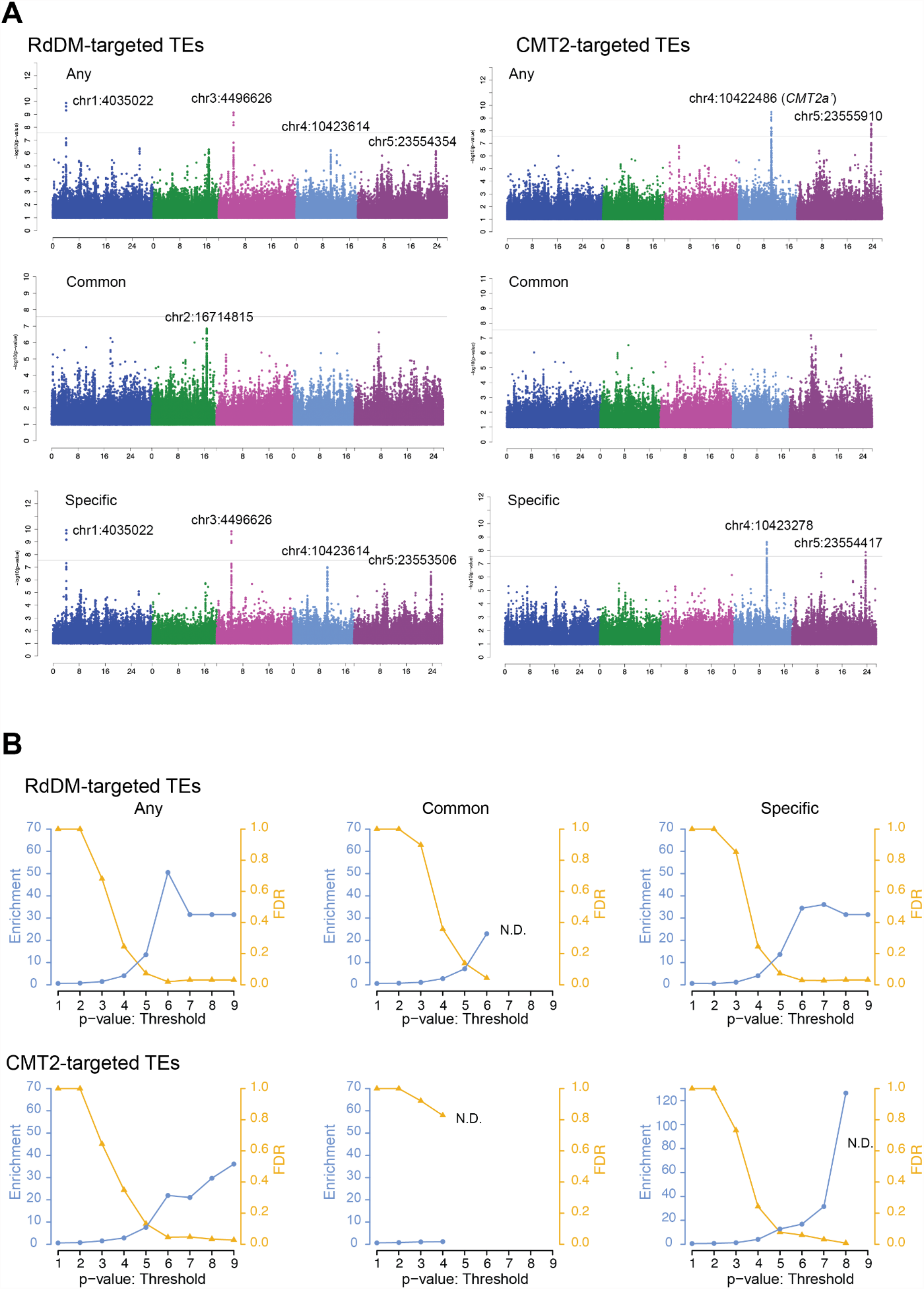
The genetic basis of mCHG and mCHH analyzed by MTMM. **(A)** Manhattan plots for any, common, specific SNP effects on mCHG and mCHH in RdDM and CMT2-targeted transposons (see Methods). Vertical lines correspond to genome-wide significance (*p*=0.05 by Bonferroni correction). **(B)** Enrichment of *a priori* genes and FDR for each GWAS result.

**Fig S13.**
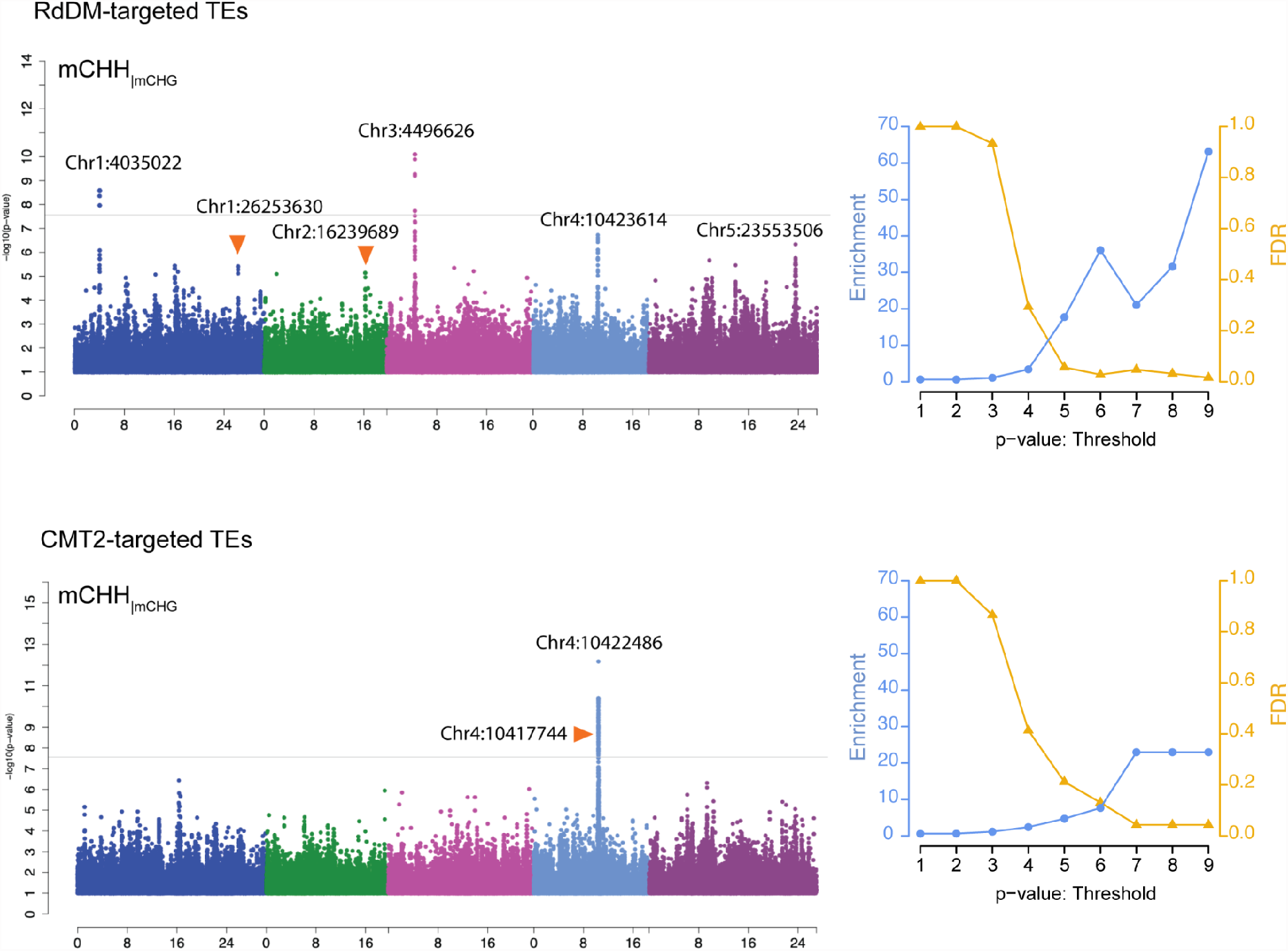
Conditional GWAS for mCHH_|mCHG_. The genetic effects on mCHH in RdDM-and CMT2-targeted transposons were analyzed by the conditional GWAS model with mCHG as cofactor. Vertical lines correspond to genome-wide significance (*p*=0.05 by Bonferroni correction). Orange arrows indicate peaks reported in previous studies as affecting mCHH (Kawakatsu et al., 2016). Each GWAS result was assessed by enrichment of *a priori* genes and FDR.

